# Discriminative Site-Directed Protein Engineering via Lightweight CASPE Platform

**DOI:** 10.64898/2026.04.24.720551

**Authors:** Qiufeng Deng, Jie Qiao, Chuan Wang, Xinyue Ni, Yongyao Chang, Nan Zhao, Rui Zhai, Haiyang Cui, Xiujuan Li, Mingjie Jin

**Affiliations:** School of Environmental and Biological Engineering, Nanjing University of Science and Technology, 200 Xiaolingwei Street, Nanjing 210094, China; State Key Laboratory of Microbial Technology, School of Food Science and Pharmaceutical Engineering, Nanjing Normal University, Nanjing, 210023, China; School of Electronic and Optical Engineering, Nanjing University of Science and Technology, 200 Xiaolingwei Street, Nanjing 210094, China; College of Life Sciences, Nanjing Normal University, No. 1 Wenyuan Road, Nanjing 210023, China

**Author notes:** Corresponding author. (M.J.), (X.L.). These authors contributed equally (Qiufeng Deng, Jie Qiao).

## Abstract

Protein large language models (PLMs) provide a novel computational paradigm for the deep mining of sequence co-evolutionary information, significantly accelerating the generation of functional proteins for biotechnological and medical applications. However, the misalignment between zero-shot predicted evolutionary fitness and industrial application requirements leads to a limited success rate in acquiring beneficial mutations, while the high training cost presents another drawback of large models. Here, we developed CASPE (Critical Amino acids Streamline Protein Evolution), a lightweight protein engineering platform for the precise localization and adaptation of critical residues, consisting of the CAS (Critical amino acid sites) and APCNet (Amino acid Point Cloud Classification Network). CAS utilizes gradient activation mapping and multi-layer attention matrices to directly extract key information determining target properties from PLMs and transform it into explicit site importance indicators, without relying on additional structural information or prior knowledge. Working in tandem with APCNet, CASPE establishes a workflow encompassing the entire trajectory from site localization to residue prediction. CASPE achieved remarkable hit rates in identifying beneficial variants for thermostability (31.3-60%) and pH tolerance (40-80%), further uncovering the potential mechanisms of action at these key target sites. Directed evolution of phytase further validated the generalizability of CASPE. CASPET-Phytase achieved a 33.3% success rate in obtaining beneficial mutants, which was significantly better than FoldX (6.7%) and ESM2-t33 (13.3%). CASPE guides enzyme evolution towards precise, site-targeted optimization, providing an efficient computational framework for developing industrial enzymes.

## Introduction

Proteins play an indispensable role in biological reactions, participating in key processes such as enzyme catalysis^1^, signal transduction^2^, and immune response^3^, and their demand in industrial applications continues to grow. However, most wild-type (WT) proteins directly derived from natural organisms exhibit limited physicochemical properties, such as thermostability and pH tolerance, rendering them difficult to meet industrial conditions directly and thus imperatively requiring engineering. Since the functional diversity of proteins is directly encoded within their amino acid sequences, identifying and optimizing functionally critical sites is of paramount importance for protein engineering. However, the potential sequence design space is astronomically vast, and functional sequences are highly sparse. Given the impossibility of an exhaustive search, researchers have developed various strategies to reduce the search space to a manageable size. The two most prevalent approaches among these are laboratory-based directed evolution and (semi-)rational design driven by computational biophysical tools (including protein structure, folding, and interactions). Although (semi-)rational design can leverage computational tools to restrict the search to a localized mutational space, it is constrained by its heavy reliance on prior knowledge, time-consuming analyses and poor generalization. Crucially, such localized exploration strategies are highly susceptible to overlooking distal allosteric networks that govern enzyme activity^4,5^, making it difficult to reconcile a global perspective with screening throughput during full-sequence scanning.

Deep learning serves as a powerful paradigm to navigate the functional protein space due to its unique advantages in data processing and feature extraction^6,7^, and it is widely applied to optimize the existing functions of natural proteins and artificially create novel functions. Among them, self-supervised PLMs provide a zero-shot mutation effect prediction method, successfully reconciling the capture of distal effects with computational efficiency^8^. PLMs capture the implicit fitness constraints of natural protein sequences from large-scale protein data, however, fitness in natural evolution is governed by a myriad of complex factors^9^, and natural sequences only impose weak-positive constraints on protein function^10,11^. It has been reported that the fitness score is biased by homologous sequences, and a too low or too high fitness score would be harmful^12^. Consequently, relying solely on the zero-shot predictions of PLMs is insufficient to substantially improve the hit rate of identifying beneficial mutants, nor can it readily generalize to different protein families and novel contexts such as industrial applications^10^.

Integrating multiple design strategies enables the more precise identification of functionally critical sites. For instance, initial designs generated by PLMs can be iteratively optimized via biophysical constraints^13^ or molecular dynamics (MD) ^14^. Although machine learning approaches that incorporate biophysical stability models yield higher accuracy, the limited scope and substantial computational expense of biophysical and MD calculations undermine the inherent speed advantage of deep learning. Researchers have proposed a directed evolutionary framework that combines PLMs with few-shot active learning, demonstrating superior predictive performance compared to zero-shot^11^. For example, EVOLVEpro leverages PLMs to extract high-dimensional protein embeddings, feeding these features alongside limited experimental data into a lightweight random forest regression model. Through iterative cycles of wet-lab validations and model updates, it enables the rapid prediction of highly active variants^10^. However, EVOLVEpro collapses the entire protein sequence into a single mean embedding vector, a strategy that inherently assumes all residues contribute equally to protein function. For enzymes characterized by high site heterogeneity, this approach risks losing critical function-related signals. Consequently, it is imperative to develop novel strategies to more precisely and efficiently identify and accommodate functionally critical sites in proteins, thereby enhancing the efficacy and transferability of AI-guided protein engineering.

To address these challenges, we developed a lightweight protein evolution platform termed CASPE (Critical Amino acids Streamline Protein Evolution, Fig. 1a-c). CASPE directly captures function-related sequence features from PLM-based high-dimensional protein embeddings, bypassing computationally intensive biophysical constraints (such as ΔG or *T_m_*). By fine-tuning functional classification models on the sequences to establish the mapping between sequences and target properties, CASPE successfully disentangles the task-specific fitness of directed evolution from the natural evolutionary fitness learned by PLMs. CASPE is a lightweight platform incorporating CAS (Critical amino acid sites) for locating critical sites and APCNet (Amino acid Point Cloud Classification Network) for predicting optimal residues. By combining Grad-CAM (Gradient-weighted Class Activation Map) ^15^ and multi-layer attention matrices ^16^ to extract the mapping relationships between amino acid sites and protein properties, CAS successfully transformed the implicitly learned protein representations into explicit site importance metrics. Moreover, the optimal amino acid was predicted by APCNet based on the microscopic three-dimensional structure and physicochemical properties surrounding the mutation site. CASPE effectively achieved the targeted modification of protein properties across dimensions such as thermostability and pH tolerance. The highest probabilities of obtaining thermostability and pH stability beneficial mutants guided by CASPE were 60% and 80% respectively. CASPE provides an effective computational framework that minimizes traditional empirical trial-and-error, enabling the precise identification of mutation sites for industrial enzyme engineering.

**Fig. 1.**
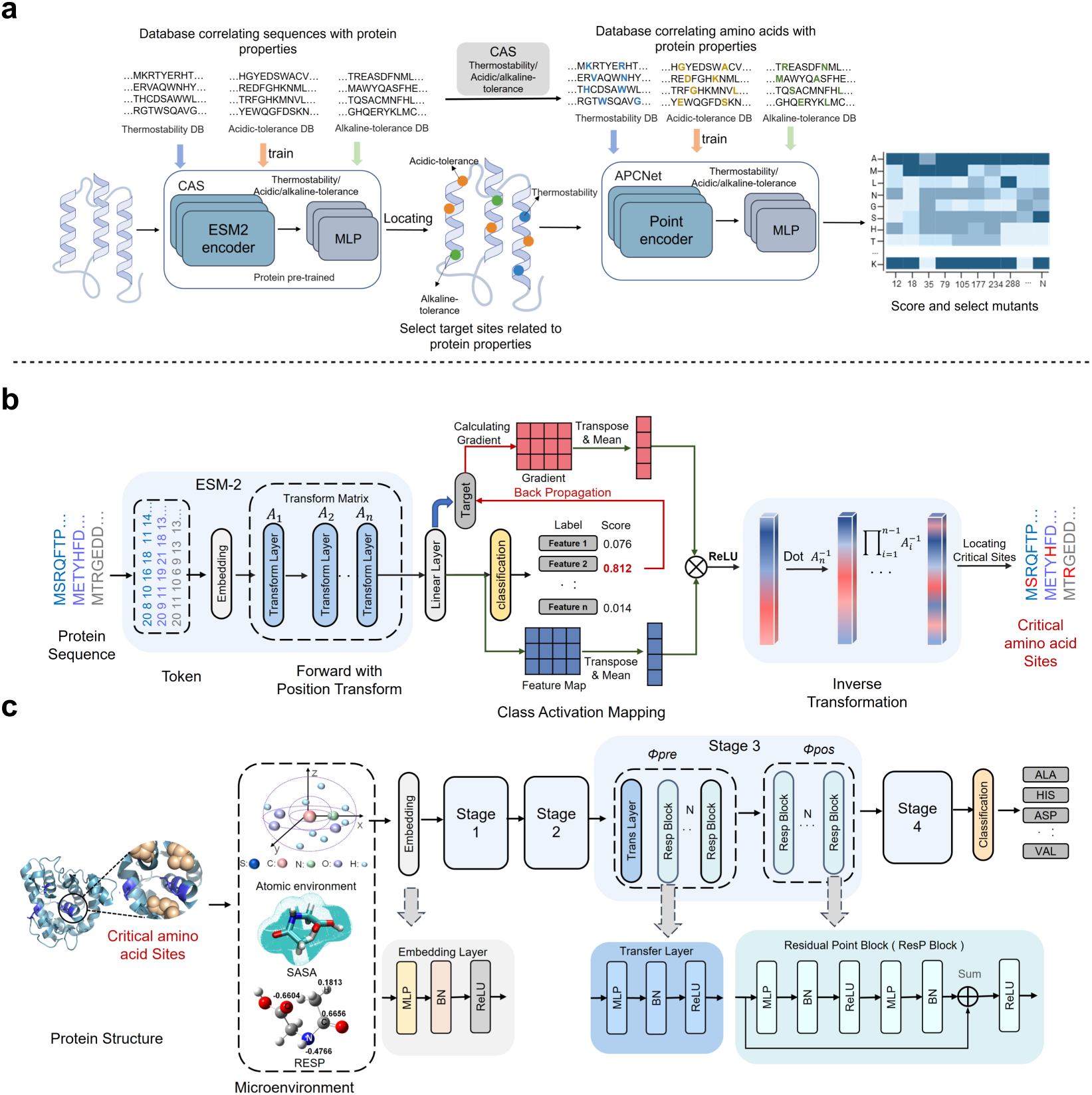
Schematic overview of the CASPE framework for precision site-directed protein engineering. **(a)** CASPE: proposed strategy for guiding the protein evolution. DB: database. CAS: Critical amino acid sites. APCNet: Amino acid Point Cloud Classification Network. **(b)** Overview of CAS: The process of locating critical amino acids via CAS by fine-tuning ESM2. **(c)** Overview of APCNet: The process of generating amino acid microenvironments, training the APCNet based on pointMLP, the stage 1-4 is the same block.

## Results

### The architecture of CASPET

We established the CASPET (CASPE for thermostability evolution) platform to drive the shift from natural evolutionary fitness to engineering-oriented single fitness via fine-tuning, enabling efficient screening of industrial enzymes (Fig. 1). For the construction of amino acid classification model, we fine-tuned ESM2, a powerful protein-related model for protein feature extraction (Fig. 1a-b and Fig. S2). The target protein sequences related to thermostability were fed into the model. To optimize the integration of ESM2, we explored two strategies (Fig. S3) across two pre-trained model sizes (ESM2-35M and ESM2-150M). Strategy 1 used ESM2 to generate training embeddings for a CNN classifier, while Strategy 2 integrated a classification layer directly onto ESM2 to eliminate embedding file storage. Since the 35M and 150M models delivered highly comparable results (Fig. S4), we proceeded with ESM2-35M. While both strategies achieved similar final accuracies across four benchmarking datasets (Figs. S5-S7), strategy 2 was ultimately selected due to its superior efficiency in identifying and locating critical amino acids.

To decipher the sequence features driving protein thermostability, we utilized CAS to extract critical sites directly from the language model’s predictive landscape (Fig. 1b and Fig. S2). As previously reported in computer vision literature, Grad-CAM exhibits inherent limitations in localizing critical features, primarily due to its difficulty in precisely delineating the boundaries of complex objects^17,18^. To account for the coarse spatial resolution of Grad-CAM, residues adjacent to the critical sites were also incorporated into the training set. To further investigate the relationship between specific amino acid compositions and protein thermostability, we calculated the relative frequencies of each amino acid across datasets before and after applying CAS (Fig. S9). The proportions of Ala and Leu had a dramatic increase after CAS, which means these amino acids have a strong correlation with the thermostability of proteins. From mesophiles to thermophiles, in the helix, the frequency of G/F/V to A/L/A is higher^19^, which indicates that Ala and Leu play critical roles in enhancing protein thermostability. The amino acid microenvironments surrounding these CAS-extracted critical sites (Fig. S10) were utilized to train our downstream screening platform, APCNet. Thanks to the rigorous data filtering provided by CAS, APCNet achieved a strong correlation with protein thermostability (Fig. 1c and Fig. S2). The training results of APCNet are showed in Fig. S11 and 12.

The extraction of critical amino acids via CAS offers two major advantages. Computationally, isolating the data most relevant to protein thermostability for downstream model training significantly alleviates computational burden and boosts predictive accuracy. Algorithmically, CAS decouples functional signals from entire protein sequences, directing the model’s attention toward critical structural microenvironments. Relying on full-length sequences often causes models to overfit to sequence homology, yielding biased predictions on similar test sets. In contrast, training APCNet on CAS-derived microenvironments prevents this bias. Theoretically, it ensures the model captures the fundamental physical rules governing thermostability rather than superficial global sequence patterns, which is highly advantageous for directed protein evolution.

### Wet-lab experimental test based on CASPET

To enhance the thermostability of cellulases using CASPET, we first employed CAS to identify 39, 24, and 37 critical mutation sites for BG, EG, and CBHI, respectively (Fig. 2g, i, k). The amino acid microenvironments of these sites were then utilized as inputs for APCNet to predict the optimal residue substitutions. Ultimately, we selected 10, 14, and 16 candidate variants for BG, EG, and CBHI, respectively, for experimental validation (Fig. 2a, c, e, and Fig. S13). Following expression and activity assays of these variants alongside their WT counterparts, we achieved hit rates of 60%, 42.9%, and 31.3% in yielding beneficial thermostable mutants for BG, EG, and CBHI, respectively (Fig. 2b/d/f, h/j/l and Fig. S14). In traditional directed evolution, the hit rate for beneficial mutants typically hovers around 1%, with the vast majority of variants being either unchanged (50-70%) or unbeneficial (30-50%)^20,21^. In contrast, the CASPE platform substantially overcomes this bottleneck, delivering a remarkably higher efficiency in identifying thermostability variants. Specifically, compared to the WT, the thermostability of BG mutants (N227A, H229F, D240N, D306A, Q319A) and EG mutants (A45N, T91I, G92S, S149N, V150L, A234T) surged by up to 205.6% and 101.9%, respectively (with specific increases detailed as 44.7–205.6% for BG and 58.8–101.9% for EG). To explore potential cross-resistance, these highly stable variants were subjected to organic solvent assays. Notably, the BG mutant D240N and four EG mutants (T91I, G92S, S149N, and A234T) outperformed the WT in the presence of 40-60% methanol, ethanol, and DMSO (Fig. S15). These robust multi-stress phenotypes indicate that mutants identified by CASPET possess untapped potential for demanding industrial applications.

**Fig. 2.**
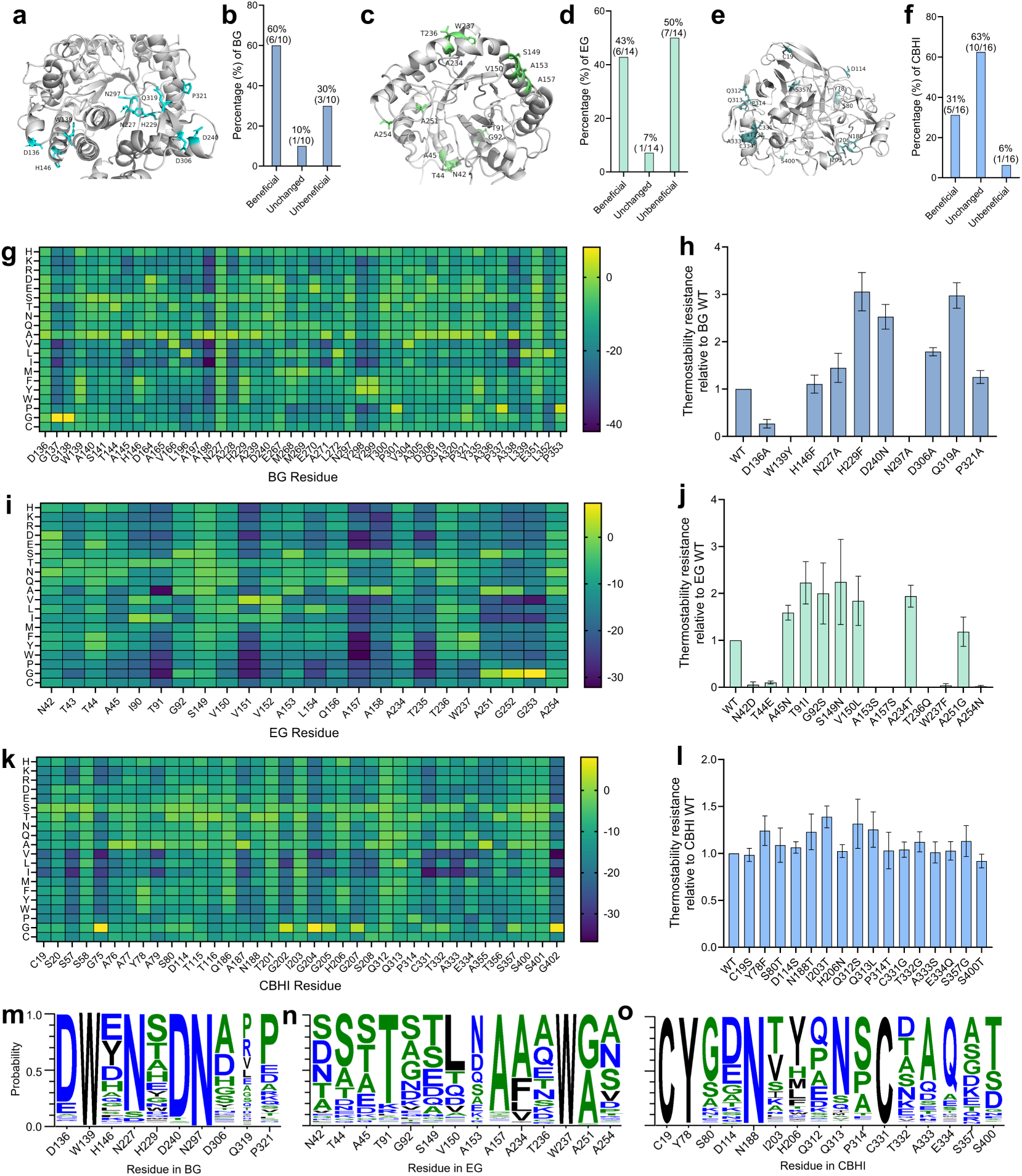
Overview of selecting single-site mutants and performance of mutants predicted by CASPET. (a, b, g, **h)** Results for BG, **(c, d, i, j)** results for EG, **(e, f, k, l)** results for CBHI. **(a, c, e)** Structures and mutation sites selected by CASPET of BG, EG and CBHI. **(g, i, k)** Heatmaps of single-site mutants selected by CASPET. **(b/d/f, h/j/l)** Performances of BG, EG and CBHI mutants selected by CASPET. **(m-o)** Conservation analysis of residues (obtained from WebLogo3 analysis).

Conservative analysis of the sites selected by CASPET was conducted (Fig. 2m-o). The amino acid conservation of some beneficial mutants of enzymes (N227A, D240N, P321A, T91I and A234T) became lower after modification. The conservation of WT enzymes may be limited by multifunctional needs (catalysis, substrate identification, allosteric regulation), while modification may simplify functions, allowing the sacrifice of partial conservation in exchange for stability^20,22,23^. Moreover, across all the sites selected by CAS, a residue’s evolutionary conservation did not strictly dictate its potential for modification (Fig. S16). Furthermore, given the limited spatial resolution of Grad-CAM, incorporating residues adjacent to the primary critical sites is essential for maximizing the hit rate of beneficial mutations. Indeed, among the successful variants identified, only T91I, V150L, Y78F, and Q313L were directly pinpointed by CAS, while the majority emerged from these adjacent positions. This outcome underscores the necessity of spatially expanded sampling to compensate for the imprecision of CAS, thereby powerfully validating the efficacy of our site selection strategy. CASPET substantially elevates the hit rate of beneficial mutants, firmly validating the efficacy of CAS in pinpointing thermostability-associated residues.

### Verification the generalizability of CASPE

#### CASPE related to pH stability

Building upon the robust performance demonstrated in thermostability prediction, we further evaluated the generalizability of CASPE by expanding its scope to pH stability. Concurrently, we collected protein sequences related to acid/alkaline stability to develop and validate this pH-targeted framework (Fig. 3a and Fig. S17). Based on CASPEA (CASPE for pH stability evolution), we identified 26 and 37 critical mutation sites in BG governing acid and alkaline stability, respectively (Fig. 3b, c). From these sites, ten candidates per condition were selected for subsequent in vitro validation (Fig. 3d-g). Notably, three mutations (D14G, A266G, and E351D) overlapped between the two sets. Ultimately, the hit rates for yielding beneficial mutants in acid and alkaline environments reached 80% and 40%, respectively (Fig. 3d-g, Fig. S18), powerfully demonstrating the broad applicability of CASPE. Relative to WT, several variants exhibited enhanced acid tolerance (D14G, T120Q, D240N, A266G, and E267Q showing 41.3-131.1% increases) and alkaline resistance (D68E, L241E, and E351D showing 23.1-80.1% increases). Notably, despite the presence of overlapping candidates during the initial computational screening, there was no overlap among the successfully validated beneficial mutants for acid and alkaline conditions. This divergence aligns perfectly with the intrinsically distinct biochemical mechanisms governing acid and alkaline resistance, confirming that CAS-identified sites are highly specific to the targeted property. Collectively, these findings establish that CASPE is not only highly effective for engineering thermostability but also readily adaptable for pH stability evolution.

**Fig. 3.**
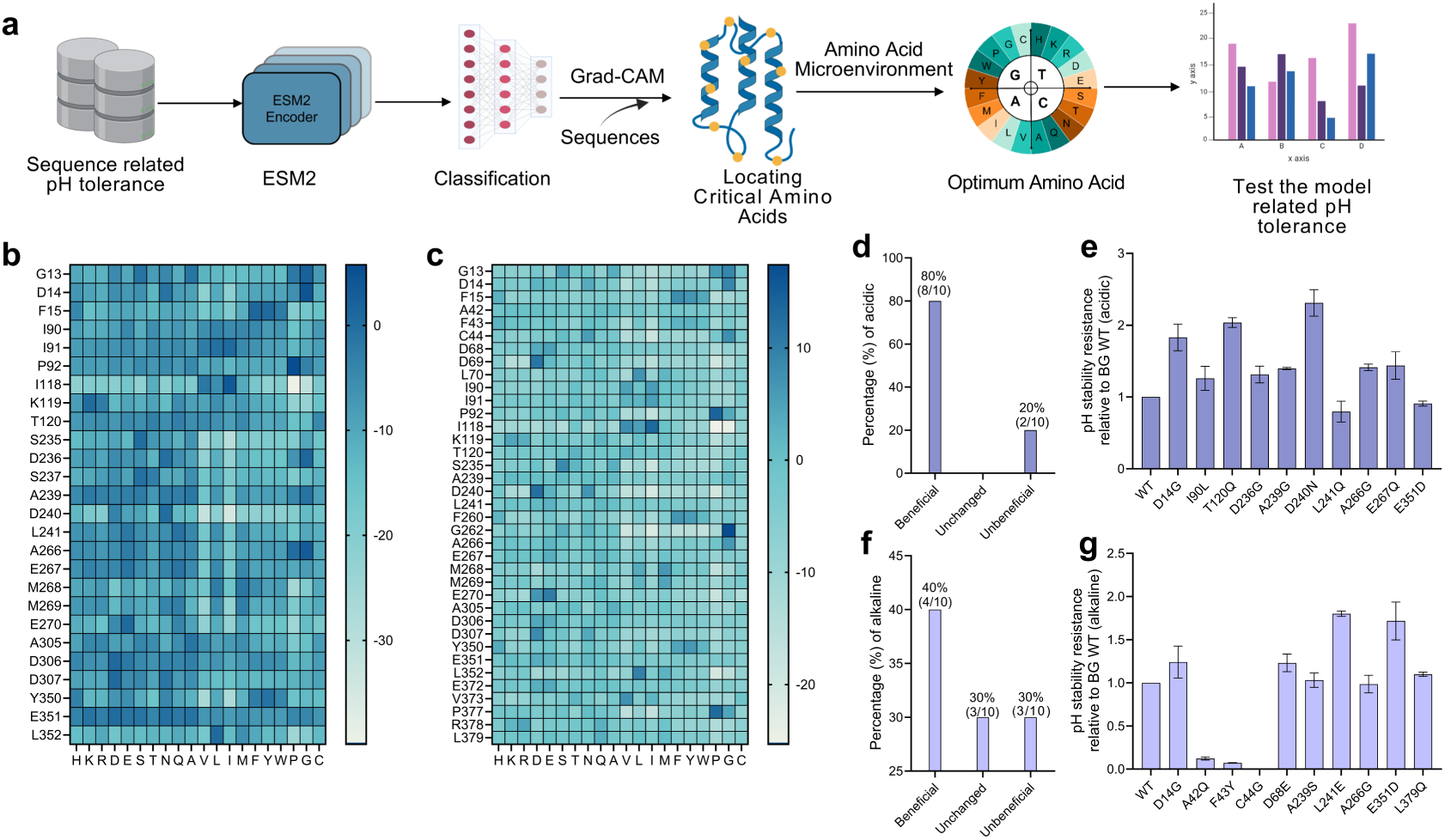
The verification of CASPE’s generalizability based on sequence related to pH stability. **(a)** The experimental logic diagram of this part. **(b, d, e)** Results for the verification of CASPE’s generalizability based on protein sequences related to acid stability. **(c, f, g)** Results for the verification of CASPE’s generalizability based on protein sequences related to alkaline stability. **(b, c)** Heatmaps of single-site mutants selected by CASPEA. **(d-g)** Performances of mutants selected by CASPEA based on BG.

To elucidate the compositional shifts driven by our model, we calculated the relative changes in amino acid frequencies (Δ proportion) pre- and post-CAS (Fig. S19). For acid stability, specific residues (K, D, E and Q) were markedly enriched, exhibiting substantial increases of 100.7%, 93.3%, 97.7% and 113.1%, respectively. A parallel trend emerged for alkaline stability, where these same residues increased by 60.1% to 159.8% (Fig. S19). Asp, Glu and Lys are acidic/basic amino acid, which could alleviate the damage to enzymes caused by extreme environments (high/low pH) by controlling the concentration of hydrogen ion. Typically, pepsin, with 13.1% of acidic residues, whose active center is formed by Asp32 and Asp215, enables its high catalytic activity at extremely low pH (∼2.0) ^24,25^. Moreover, salt bridges are helpful for enhancing the stability of proteins ^26–28^, and D, E and K are the critical amino acids in salt bridges^29^. K, D, E and Q are hydrophilic amino acids, which could bind water molecules to form a hydrated shell, protecting enzymes from denaturation caused by high/low pH^30^. Furthermore, except for the E351D variant, the beneficial mutants generally exhibited lower evolutionary conservation (Fig. S20). This observation aligns with the established paradigm that thermostability is often enhanced at the expense of evolutionary conservation. Consistently, APCNet-predicted sites showed no strict correlation with residue conservation (Fig. S21), further validating that our model prioritizes underlying biophysical rules over historical evolutionary constraints.

#### Finetuning CASPE based on phytase sequences

Our results indicate that the effectiveness of general models in identifying beneficial mutants varies across different enzymes (Fig. 2h, j, l). To fill this gap, we demonstrated that training protein-specific models significantly improves the efficiency of enzyme evolution. Phytase sequences obtained from three databases were used as the input of CASPE for data screening (Fig. 4a and Fig. S22). CASPEA-Phytase (CASPEA for phytase) was developed based on phytase sequences for improving phytase pH stability (Fig. S23). 15 mutants based on CASPEA and CASPEA-Phytase were selected for wet-lab experiments (Fig. S24). Our results indicate that the baseline success rate for identifying beneficial mutants using the general CASPEA model was relatively low (Fig. 4b, e, and Fig. S25). However, by employing the customized CASPEA-Phytase model, the yield of beneficial mutants with improved acid stability significantly increased from 13.3% to 66.7%, while the proportion of alkaline-stable mutants rose from 40% to 53.3% (Fig. 4c, d, f, g, and Fig. S26). WT phytases typically exhibit minimal activity at high pH ^31^ and researchers have tried to improve its alkaline stability. A recent state-of-the-art automated platform reported a hit rate of 55% for beneficial phytase mutants, but this required the construction and screening of a substantial library (180 variants) in the initial round^32^. In our study, a comparable success rate (53%) was achieved while reducing the mutational search space and wet-lab workload.

**Fig. 4.**
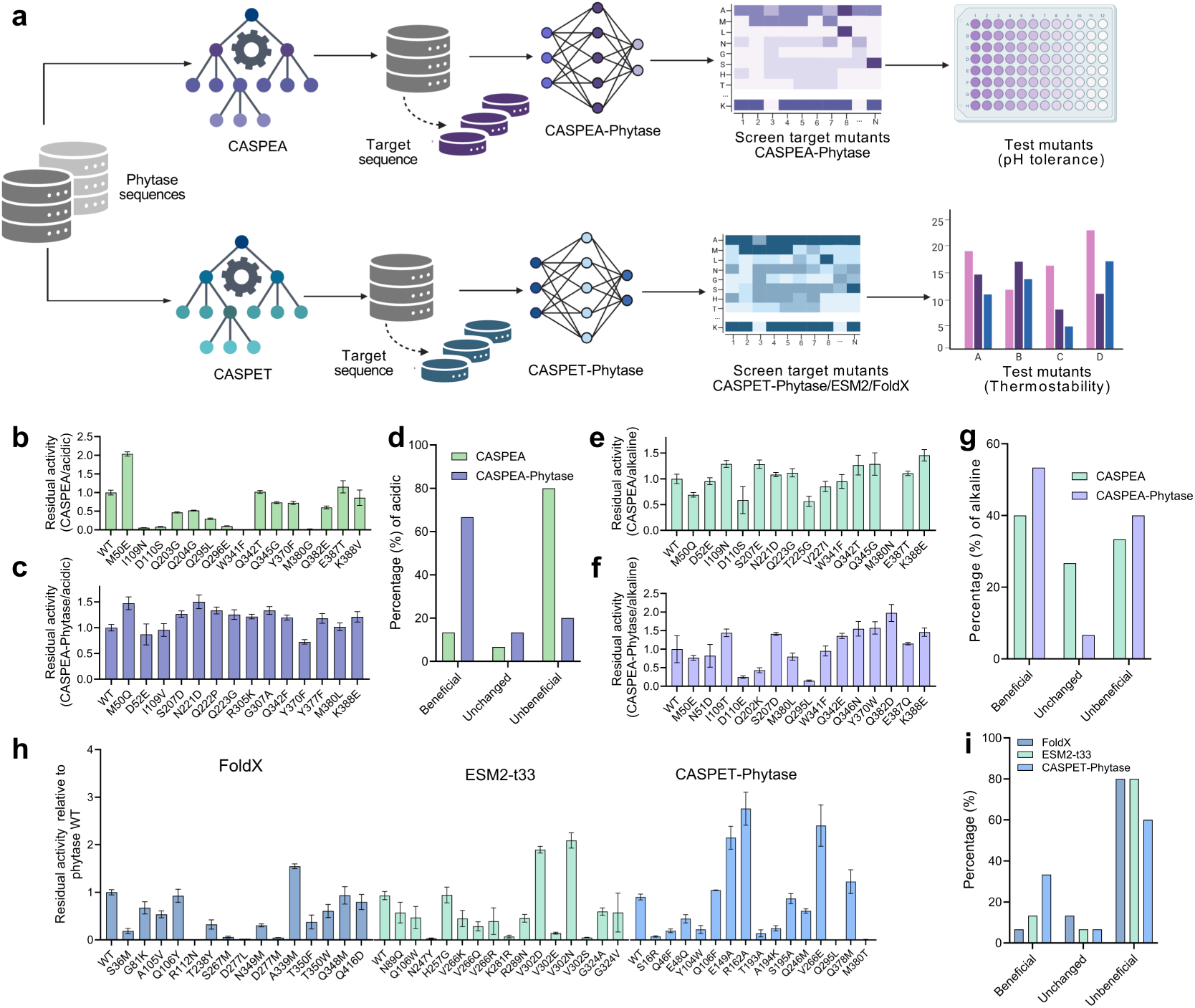
The verification of CASPE’s generalizability based on phytase sequences. **(a)** The experimental logic diagram of this part. **(b, e)** Performances of mutants selected by CASPEA. The model is trained on protein sequences related to acid/alkaline stability (the same with Fig. 3). **(c, f)** Performances of mutants selected by CASPEA-Phytase. The model is trained on phytase protein sequences related to acidic/alkaline stability. **(d, g)** Summary of mutants predicted by general models or specific models. **(h, i)** Performances of mutants selected by FoldX, ESM2-t33 and CASPET-Phytase.

To evaluate the performance of the CASPET-Phytase (CASPET for phytase), we compared its efficiency in identifying beneficial mutants against that of FoldX and ESM2-t33. This was achieved by experimentally testing the top 15 mutant candidates predicted by each method. The specific candidates for FoldX, ESM2-t33, and CASPET-Phytase are detailed in Figs. S28-S30. CASPET-Phytase identified beneficial mutants with significantly higher thermostability gains (e.g., R162A, increased by 206%) compared to the best variants from FoldX (A339M, +54.4%) and ESM2-t33 (V302N, +124%) (Fig. 4h, Fig. S31). Furthermore, CASPET-Phytase demonstrated superior screening efficiency, achieving a 33.3% success rate, which substantially outperformed ESM2-t33 (13.3%) and FoldX (6.7%) (Fig. 4i). Furthermore, the 12.5M-parameter CASPET-Phytase model is exceptionally lightweight compared to ESM2-t33 (650M). Unlike widely used tools (e.g., FoldX and ESM2) that depend on computationally intensive full-sequence saturation mutagenesis to screen for functional variants^33–35^, our platform leverages CAS to precisely pinpoint critical mutation sites. Consequently, this targeted approach confirms that CASPE is a highly efficient and computationally economical method for protein evolution. Conservation analysis of the mutation sites screened by the FoldX, ESM2-t33, and CASPET-Phytase indicates that the mutants identified by FoldX and CASPET-Phytase exhibited no significant correlation with amino acid conservation (Fig. S32a, c), and the conclusion for CASPET-Phytase is consistent with the previous discussions. As mentioned before, fitness calculations in large language models are inherently biased by sequence homology. Consistent with this observation, the variants prioritized by ESM2-t33 exhibited increased evolutionary conservation. For instance, at positions V302 and V266, the model-identified substitutions (V302D/N/E/S and V266Q/R/K) all introduce amino acids that display higher conservation levels than the WT residues at their respective sites (Fig. S32b).

### CASPE evaluation *in silico* and interpretability analysis of APCNet

To further elucidate the APCNet model, we first employed CAS to screen for critical sites within protein sequences from the ProteinGym^36^ and CDNA^37^ databases. We then evaluated the fitness prediction performance of APCNet on these CAS-filtered datasets. We extracted 10 amino acids that were the most relevant to the thermostability of proteins by CAS. Spearman and Kendalltau correlation coefficients reflect the rank relationship between two sets of data, while Person correlation coefficient reflects the degree of association between the two sets of data. Specifically, APCNet outperformed ESM1b^38^, ESM1v^8^, ESM2^6^, GEMME^39^, MSA1b^7^, VESPAI^40^, CARP-640M^41^, MULAN-middle^42^, RITA-large^43^, Progen2-base^44^, Tranception-L^45^ and EVmutation^46^ (Fig. 5a, b and Fig. S33-35). While existing large models possess remarkable predictive capabilities, APCNet offers a highly compact alternative with only about 12.5 million parameters. Notably, by leveraging CAS for initial site screening, this lightweight model achieves highly competitive in some specific cases. This suggests that CAS is an effective and computationally efficient tool for pinpointing critical amino acid sites associated with protein thermostability. We performed PCA on thermostability-related sequences from ProteinGym and CDNA to evaluate feature extraction. Remarkably, CAS-selected data concentrated more informational variance into PC1 (35.1%) compared to both the full sequence (33.3%) and random selection (32.5%). Furthermore, CAS mitigated the dispersion of variance into PC2 and PC5, lowering them to 19.8% and 5.8% (down from the 20.0% and 6.2% baselines), thereby avoiding the variance inflation observed with random selection (21.3% and 6.4%) (Fig. 5c-e, Fig. S36). These dimensional redistributions confirm that CAS effectively filters noise and selectively targets amino acid residues crucial for thermostability. Additional details are available in Supplementary Text 1.

**Fig. 5.**
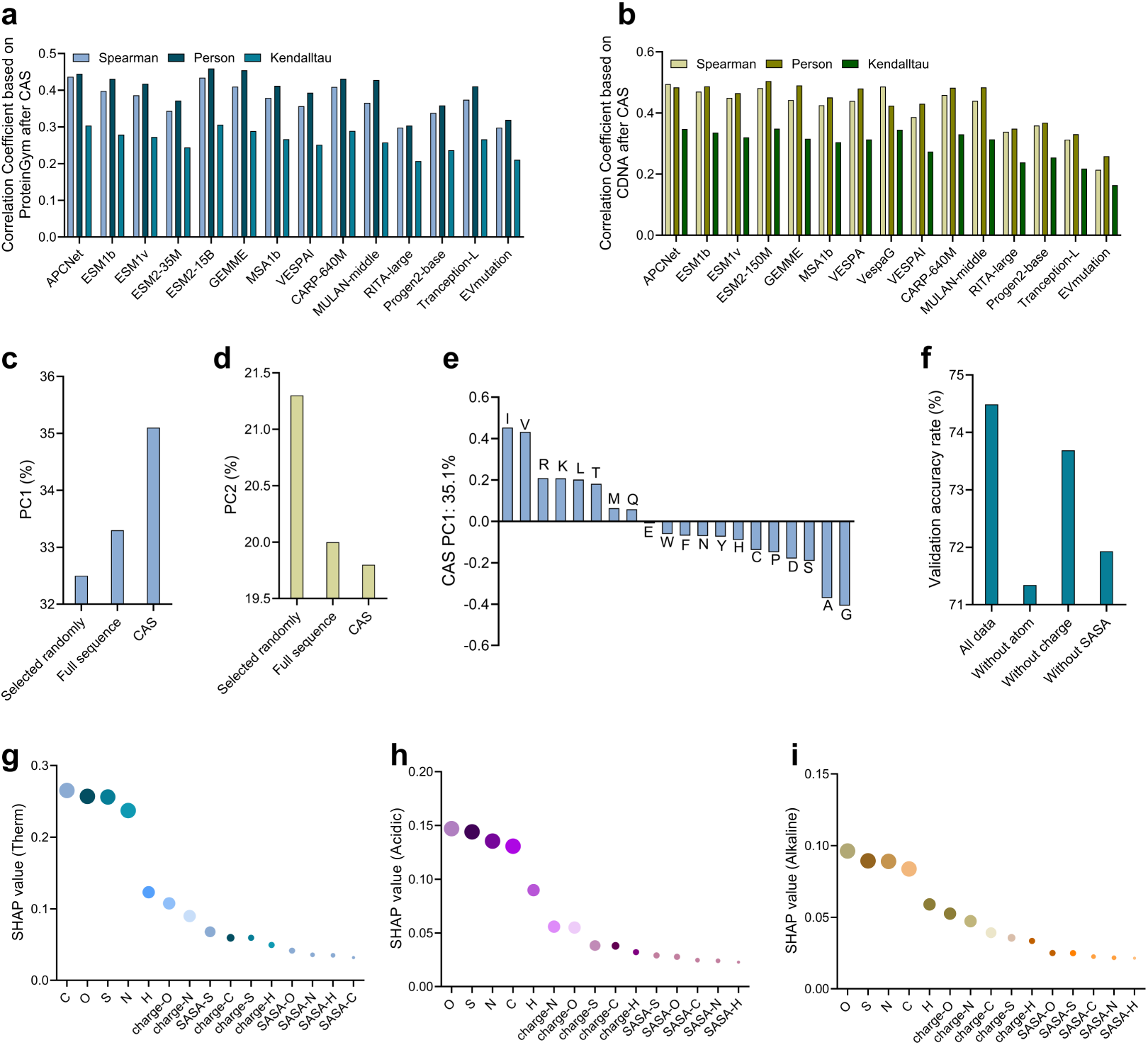
CASPE test *in silico* and interpretability analysis of APCNet. (a, **b)** the results of correlation coefficient analysis based on proteinGym **(a)** and based on CDNA **(b)**. **(c-e)** results of PCA analysis. **(c, d)** PC1-2 of CAS, full sequence and data selected randomly. CAS: 10 sites were selected from each sequence by CAS. Data selected randomly: 10 sites were randomly selected from each sequence. **(e)** PC1 of CAS. **(f)** The ablation experiment results, there are three types of amino acid microenvironments, the accuracy changes of models were observed by removing atoms, charges or SASA respectively. **(g-i)** Global analysis of each feature based on SHAP, the influence of each feature is measured by calculating the average absolute value of each feature in all samples. **(g)** for thermostability, **(h)** for acid stability, **(i)** for alkaline stability.

Having established that the CAS strategy effectively concentrates thermostability-relevant information and enables APCNet to achieve superior predictive performance, we next sought to decode the underlying biophysical rules driving the model’s decisions. To demystify how APCNet evaluates specific structural features, we adopted three interpretability tools, including SHAP, Captum, and Permutation Importance analysis, to evaluate the importance of three major amino acid microenvironments (atoms, atomic charges, and SASA). Cross-validation using SHAP, Captum and Permutation Importance revealed a highly consistent global hierarchy in feature reliance, uniformly ranking the macro-contributions of atoms first, followed by charges and SASAs. While the exact ranking of the five individual atom types exhibited slight algorithmic-dependent variations due to their distinct attribution mechanisms (Fig.5 g-i and Fig. S37-40). Furthermore, an analysis of 15 local features across all 20 amino acids revealed a shared global trend in both the thermal and pH stability models (Fig. S41). Both SHAP and Captum analyses proved that APCNet always gives the highest priority to atoms, followed by charges and SASAs. Both the global and residue-specific analyses point to the same conclusion, demonstrating that this overarching trend remains identical across all evaluation scales. Nevertheless, the precise rankings of the 15 sub-features show noticeable variations between the thermostability and pH stability. This demonstrates that while the core predictive logic of APCNet is grounded in universal physical rules, the model retains the flexibility to finetune its micro-environmental parameters according to the specific properties of proteins. To complement the interpretability analysis, we performed ablation experiments by removing atoms, charges, or SASA from the amino acid microenvironments (Fig. 5f and Fig. S42). Interestingly, while the above methods positioned charges ahead of SASAs, the ablation study flipped this order. This divergence highlights the distinction between the two approaches, where attribution tools measure the immediate predictive contribution of a feature, whereas ablation studies evaluate the model’s global vulnerability and redundancy when a feature is completely lost.

## Discussion

PLMs effectively capture general fitness in natural evolution, which limits their industrial applications. Here, we developed CASPE, a deep learning-guided framework that explicitly decodes the implicit representations of protein language models into quantifiable functional metrics, which successfully translates the natural evolutionary fitness into specific target traits for directed evolution. By disentangling this fitness landscape, we highlight that natural evolutionary fitness is not entirely synonymous with survival under the extreme processing conditions of the industry, such as extreme pH and high temperatures. Cagiada et al. demonstrated that explicitly deconvolving biophysical constraints from evolutionary priors provides a direct route to precisely pinpoint high-value mutational targets ^47^. Recent study has deconvolved the evolutionary probabilities predicted by large language models into function and protein stability to identify functional residues^48^. Similarly, CASPE could identify critical residue related to the specific property of proteins by decoupling task-specific fitness from general evolutionary fitness.

More broadly, this work reframes how sequence space should be evaluated to guide directed evolution. PLMs can extract complex protein information from massive natural protein sequences by masked language modeling (MLM). These models mostly adopt the *ΔLL* of masked protein sequence to evaluate zero-shot prediction of mutational effects, thereby accelerating the screening of beneficial mutants^8^. They rely on *ΔLL* to score and rank all possible mutations and select the top-*K* candidates with the highest scores^32,49–51^. The predictions of PLMs are governed by site-specific contexts, current mainstream PLMs typically employ *ΔLL* to rank mutations across the entire sequence. This practice introduces a critical limitation by implicitly assuming a homogeneous contextual influence across all sites. In bio-catalysis, protein sequences exhibit significant site heterogeneity ^52,53^, for amino acid substitutions at active center or allosteric sites contribute more to overall fitness than those at other residues^54,55^. Statistically, after decoupling sequence site-specific context differences, we deduced that *ΔLL* ranking serves as a proxy for general fitness in natural evolution. They fundamentally assess the capacity to preserve protein stability following, in which higher rankings of the predicted mutation reflect greater sequence conservation, rather than conferring directed evolutionary advantages for specific industrial applications (Supplementary Text 2). While this theoretical implication corroborates the known propensity of ESM2 to prioritize conserved residues, neither CASPE nor FoldX demonstrates a corresponding conservation bias (Fig. S32). This divergence underscores the unique value of our supervised attention-extraction framework. By unbinding the search space from evolutionary conservation, CASPE is able to navigate previously inaccessible regions of the fitness landscape, successfully identifying non-conserved mutations that dictate industrial resilience.

Translating this theoretical advantage into practical application, CASPET was highly efficient in guiding protein thermostability evolution, as evidenced by the wet-lab experiments based on cellulase (Fig. 2). We tested the generalizability of CASPE based on protein sequences related to acid stability or alkaline stability, CASPEA efficiently screened out beneficial mutants (Fig. 3). We also proved the generalizability of CASPE based on a certain type of protein (phytase). It was found that enhancing protein properties based on amino acid sites related to the property of protein would significantly improve efficiency of directed evolution compared with traditional methods (Fig. 4). By narrowing down the search space, CASPE helps alleviate the experimental burden, providing a practical and streamlined discriminative strategy for industrial bio-catalysis. CASPE effectively mitigates the homology bias between training and test sets through CAS-based data selection. Furthermore, it efficiently identifies critical target sites for subsequent protein evolution. Additionally, CASPE requires limited resources and could be trained on a single NVIDIA RTX 4090 within a few days due to its targeted data selection.

Despite the robust predictive performance of purely sequence-based representation learning, we acknowledge its inherent limitations in fully capturing drastic backbone rearrangements or highly complex local microenvironments. To address this spatial blind spot, the field is currently witnessing the emergence of multi-modal representation models that integrate sequence, three-dimensional structural, and physicochemical features. Furthermore, for highly dynamic physical processes, integrating explicit MD simulations remains an indispensable complementary strategy. Crucially, as a highly modular framework, CASPE is not strictly bound to current sequence-only PLMs, it offers plug-and-play compatibility with these emerging spatially-aware models and can seamlessly interface with future, more expressive biological foundation models.

## Methods

### Database: Protein sequence dataset

All the protein sequences related to the thermostability in this article are from the database learn2therm^56^, which contains a total of 69 million pairs of protein sequences and hosts with the optimal growth temperature of the host. The protein sequences from the hosts with the optimal growth temperature above 60 ℃ (excluding 60 ℃) were collected. Studies have shown that the OGTs (optimal growth temperatures) of the host are closely related to the optimal temperature or TM (melting temperature) of proteins^57^. There are approximately 280,000 sequences after filtering. More details about data distribution are shown in Fig. S1. In order to obtain a model with better training performance, we segmented the dataset for 4 parts as shown in Table S1: the temperature range of the dataset for Model 1 are 20-25 °C, 50-55 °C and 70-80 °C; the temperature range of the dataset of Model 2 are 32-35 °C, 55-60 °C and above 80 °C; the temperature range of the dataset of Model 3 are below 20 °C, 38-45 °C and 65-70 °C; the temperature ranges of the dataset for Model 4 are 26 °C, 45-50 °C, and 60-65 °C. The training dataset, verification dataset and test dataset are divided into 8:1:1 ratio in different temperature ranges. The protein sequences concerning pH stability were acquired based on Rong et al. ^58^ . The protein sequences of this database were obtained based on the optimal pH of microorganisms. We screened the sequence whose length was as greater than 100 and less than 1000, and obtained a total of 84,303 acid stability and 49,912 alkaline stability sequences, respectively. The training set, validation set and test set were divided respectively according to the 8:1:1 ratio of the acid resistance and alkaline resistance sequences. We obtained related sequences from the Uniprot, PDB, and NCBI through the keyword “phytase”, yielding a total of 52,295 sequences. We selected sequences based on specific requirements. In details, 6,736 sequences were identified for thermostability (optimum temperature above 50°C), 27,054 for acid stability, and 25,241 for alkaline stability.

### Constructing and fine-tuning classification models

When constructing classification model, taking advantage of the analytical ability of the existing large models for protein sequences, the transform blocks of ESM2 (Fig. 1b) was adopted as the encoder^6^ of the classification model to encode the input protein sequences to the high-latitude feature vectors (embeddings). ESM2 was connected by MLP as a classifier to classify embeddings. To prevent the overfitting of models, BatchNorm1d, ReLU, and Dropout layers were added to the classification layers. Since one dimension of embeddings was related to the length of the input protein sequence, zero padding was applied to expand it to a fixed length. The embeddings were regarded as the input of the subsequent MLP classifier after one-dimensional expansion. As for the selection of the loss function, cross-entropy was applied for the loss function of classifier and introduce L2 regularization to prevent overfitting. The calculation formula is as follows:

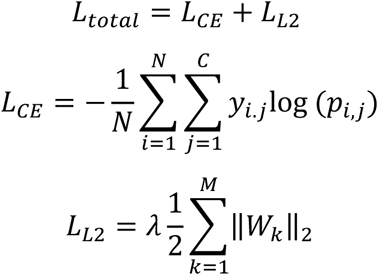

Where, N is the number of samples, C is the number of categories, 𝑦_𝑖.𝑗_ is the one-hot encoded value of sample 𝑖 for category 𝑗, and 𝑝_𝑖,𝑗_ is the prediction probability of the model that sample 𝑖 belongs to category 𝑗. 𝜆 are the regularization coefficient (hyperparameter and regularization strength), M is the number of layers of the neural network, 𝑊_𝑘_ is the weight matrix of the 𝑘^𝑡ℎ^ layer,

‖ ‖_2_ is the L2 norm of the matrix, and the final loss function 𝐿*_total_* is the sum of the cross-entropy and the L2 norm of the model weights.

We loaded the EMS2-35M weight file as the pre-training model, and the transformer encoder consists of 12 layers and 20 attention heads, with 35 million parameters. When fine-tuning the classification model, the learning rate was set to 1 × 10^-4^, the optimizer was Adam, and the epoch was 200. The learning rate gradually decreased according to the cosine function curve. When the test set loss tended to be stable, the training was manually stopped. The model was trained on a RTX 4090 24G GPU, and the time depended on the size of the dataset and the trend of loss stabilizing, which was approximately 2-11 days.

### Localization of critical amino acids

As for inferring the model, each point in the amino acid sequence had a different contribution value to the final classification weight, the larger the contribution value, the closer the correlation between the point and the characteristics of the protein (Fig. 1b). In order to locate the critical amino acids, we utilized the critical data in the model inferring process to conduct reverse inferring on them, and inferred the contribution value of each point on the amino acid sequence from the weight values of the classification. The process was mainly divided into two steps. Firstly, we obtained the contribution value of each position in the embeddings through Grad-CAM. The formula is as follows:

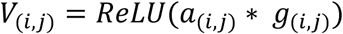

Where 𝑎_(𝑖,𝑗)_is the activation value of the 𝑖^𝑡ℎ^ row and 𝑗^𝑡ℎ^column element in embeddings when the model propagates forward to the linear layer, and 𝑔_(𝑖,𝑗)_ the gradient of the classification weights to the element of the 𝑖^𝑡ℎ^row and 𝑗^𝑡ℎ^column of embedding when the model is backpropagated to the linear layer, 𝑎_(𝑖,𝑗)_is multiplied by 𝑔_(𝑖,𝑗)_, and the ReLU function was used to filter the positions that have a positive impact on classification. Secondly, the embeddings contribution was converted to the amino acid sequence contribution as follows:

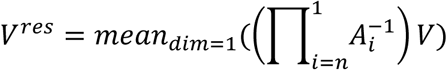

Where 𝑉^𝑟es^ is the contribution value of amino acid in the full sequence, 𝑉 is the contribution value matrix of embeddings, 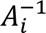 is the inverse matrix of the attention weight matrix of the 𝑖^𝑡ℎ^ transformer layer, mean_dim=1_() means to calculate the one-dimensional mean, and we use the mean contribution value of the 𝑗^𝑡ℎ^ column as the contribution of the 𝑗^𝑡ℎ^ element of the amino acid. In the encoder process of ESM2, embeddings were obtained by the attention matrix of multiple transform layers of the amino acid sequence. Therefore, the contribution values of the embeddings could be converted into the contribution values of the amino acid sequence by the inverse transformation of attention matrix (Fig. 1b and Fig. S8). We used the weight files obtained above to infer the existing protein sequences. Firstly, we selected all sequences with the OGT above 60℃ based on the classification results. Then, the amino acid contribution values were obtained through the above methods, and all amino acids in a sequence were sorted based on the contribution values to extract several amino acid sites most related to thermostability of protein. Considering the ambiguity of Grad-CAM, the critical sites and their adjacent residues were selected as the final selected sites.

### Microenvironment data sampling

There were three classes of microenvironments around amino acids, including atomic environment, SASA (solvent accessible surface area), and partial charges of atoms (Fig. 1c and Fig. S10). The atomic environment included C, H, O, N, S, and the environment of H was obtained by PDB2PQR, which could add H of PBD file^59,60^. For atomic radius, we adopted the Van der Waals radius of C, H, O, N, and S, which are 1.7Å, 1.1Å, 1.52Å, 1.55Å, and 1.8Å, respectively^61^. SASA was obtained by FreeSASA^62^, which is used to characterize the hydrophilicity and hydrophobicity of proteins. The atomic charge is RESP (Restrained Electro Static Potential), and it is currently the most suitable charge for molecular simulation of flexible small molecules^63^. Multiwfn^64,65^ was adopted to calculate the RESP charges of 20 amino acids.

In order to preserve the integrity of the sampled data to the greatest extent, we adopted a 3D (three-dimensional) Point Cloud data format to describe the microenvironment around amino acids. The main feature of 3D Point Cloud data is that it can fully preserve the original geometric information in 3D space without discretization^66^. The spatial information could be more accurately described by 3D Point Cloud data. During sampling, we constructed a three-dimensional coordinate system with the 𝐶α → 𝐶 of the central amino acid as the X-axis direction and the 𝐶α → 𝑁 as the Y-axis direction, a quantitative description of the above environments were in terms of (x, y, z, value), in the form:

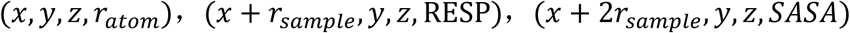

Among them, (𝑥, 𝑦, 𝑧) are the precise coordinates of the current atom, 𝑟*_atom_* is the Van der Waals radius of the atom, and 𝑟_sample_ is the sampling radius of the microenvironment. Since the spatial position is unique, the X-axis offset method was adopted to describe the remaining two characteristics. This method would not lead to the incorrect information because the offsets of the data format were uniform, and the model would automatically adapt to this type of data during training.

### Target gene selection and synthesis

Three main cellulases were selected for model validation, including CBHI, EG and BG^67^. Among them, the similarity between the CBHI sequence and 4D5Q^68^ (PDB ID) is 99.5%, and the similarity between the EG sequence and 5L9C^69^ (PDB ID) is 96.8%, BG (H0HC94, EHJ97046.1) is reported in the literature^70^. The information of the three sequences is shown in Table S2. After the synthesis of the CBHI and EG genes, they were assembled into the pBSYA1S1Z^71^ plasmid, and plasmids were transformed into *Pichia pastoris* BSYBG11 for expression. For BG, the target gene was assembled onto the pET-21b plasmid, and subsequent protein expression was carried out in o *E.coli* BL21^72^. The sequence applied for validating the phytase-based models was obtained from the literature ^32^. The target gene was synthesized and assembled onto the plasmid pET-28a plasmid, which was then expressed in BL21.

### Selection of the target mutations

After obtaining the model training results, it is necessary to select the mutation sites based on the CASPE. Specifically, we got the critical sites of the sequences that need to be modified based on protein classification models. Considering the ambiguity of the Grad-CAM, we simultaneously obtained two adjacent residues of each critical site. After removing the repeated sites, mutation sites selected by the classification models were the alternative mutation sites of the target sequence. Based on the above critical amino acid sites, 3D Point Clouds of the critical amino acids were generated, and they were regarded as the input of the APCNet to predict their optimal amino acids. We stopped to screen the mutants until the number of mutations was more than ten, which were then verified by subsequent wet-lab experiments.

### Conservative analysis of amino acid sites

After obtaining amino acid sites related to protein thermostability, acid stability or alkaline stability based on CASPE, we found all sequences with similarity between 35% and 90% ^73^ to the target sequences (CBHI, EG and BG) from NCBI and PDB databases. We got 154, 87 and 1476 sequences for CBHI, EG and BG, respectively. All sequences were processed in MEGA11^74^ to retain the sequences selected based on Grad-CAM and the sites further screened by APCNet. Subsequent amino acid conservation analysis was performed in WebLogo3^75^.

### Enzyme activity test and characteristic characterization

The activity of CBHI was detected in 96-well MTPs with the resorufin-D-cellobioside (Res-CB) as the substrate^76,77^. All enzymes (WT and variants) were placed in sodium acetate buffer (0.1M, pH 4.5), incubated at 25 °C or 45 °C (900 rpm) for 30 minutes, and the fluorescence was measured for 30 minutes by a microtiter plate reader to detect the enzyme activity. The activity of EG was tested in 96-well MTPs with the modified Azo-CMC method^71,78,79^. All enzymes (WT and variants) were incubated at 25 °C or 78 °C (900 rpm) for 30 minutes in sodium acetate buffer (0.1M, pH 4.5). Similarly, all BGs were incubated at 25 °C or 55 °C for 45 minutes, and the activity of them were detected with 4-nitrophenyl β-D-glucopyranoside (pNPG) as the substrate ^80^. For phytase, the activity of phytase was tested based on 4-MUP ^31,81^ (4-methylumbelliferyl phosphate, Sigma #M8168) and the enzyme activity was determined through the variation of fluorescence. The residual activities of mutants (at room temperature or high temperature) were expressed by dividing the mutant activity by the WT activity, as shown below:

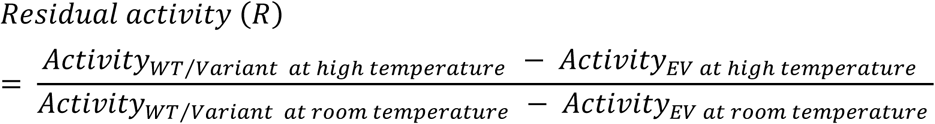

While the thermostability resistance is calculated as follows:

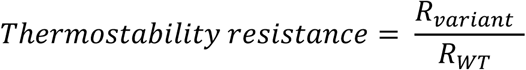

As for testing the pH stability of enzyme, we verified the transferability of CASPE on BG. Specifically, all enzymes were incubated in sodium acetate buffer with different pH (pH 4, 7 and 11) for 20 minutes or 30 minutes (pH 4 and 7 for 30 minutes, pH 7 and 11 for 20 minutes), and the activity of BG was detected as mentioned above. While all enzymes related phytase were incubated in tris-maleate buffer (0.2M) with different pH (pH 4.5 and 5.6 for 30 minutes).

The residual activity of the mutant in different pH was expressed by dividing the mutant activity by the WT activity, as shown below:

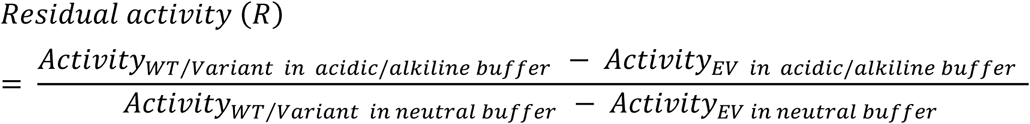

While the pH stability resistance is calculated as follows:

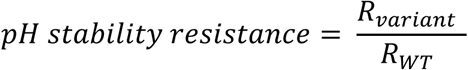

### Test CASPE with other models based on Zero-shot prediction

To test the performance of CASPE, we applied the DMS (deep mutational scanning) data from ProteinGym^36^ and CDNA^37^ to test CASPE. ProteinGym is a benchmark for predicting the fitness of large protein models, while CDNA is a database related to protein thermostability. To test the performance of CASPE, we selected the critical amino acids of target sequences from ProteinGym and CDNA based on Grad-CAM. For a specific amino acid site, the weights of 20 amino acids would be inferred by the model based on the surrounding amino acid microenvironment. The greater the weight of an amino acid, the higher the probability of its occurrence, it would be considered as the most suitable amino acid at this location. After obtaining the weights, we determined the influence of the mutant according to the following formula:

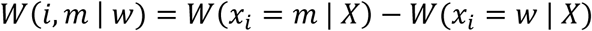

Among them, 𝑊(𝑖, 𝑚 | 𝑤) is the weight of the single mutant, 𝑊(𝑥_𝑖_ = 𝑚 | 𝑋) is the weight of the mutant, 𝑊(𝑥_𝑖_= 𝑤 | 𝑋) is the weight of the WT, 𝑖 is the 𝑖^𝑡ℎ^ amino acid, 𝑋 = (𝑥_1_, 𝑥_2_, …, 𝑥_𝐿_) is the amino acid sequence, L is the length of sequence. We calculated the correlation between zero-shot prediction of APCNet and proteinGym or CDNA based on the critical amino acids selected by Grad-CAM. Pearson, Spearman and Kendall correlation coefficient were applied to evaluate the performance of different models. We referred to the stats module of Scipy in python to calculate the above three correlation coefficients. Their calculation formulas are as follows:

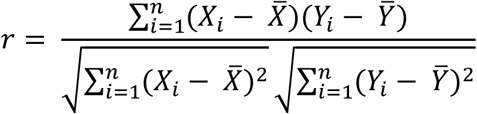

The above is the formula for calculating the Pearson correlation coefficient, where 𝑋_𝑖_ and 𝑌_𝑖_ are the variable values and *X̄* and *Ȳ* are the average value.

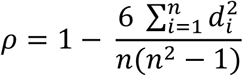

The above is the formula for calculating the Spearman correlation coefficient, where d_𝑖_ is the difference in rank between the two variables and n is the sample size.

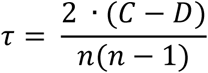

The above is the formula for calculating the Kendallta correlation coefficient, where C is the consistent logarithm (with the same sorting direction), D is the inconsistent logarithm (with the opposite sorting direction), and n is the sample size.

### PCA analysis based on Grad-CAM

PCA (Principal Component Analysis) is a commonly tool that can reduce the dimension of data to achieve the simplification of data. We conducted PCA analysis on the data from proteinGym and CDNA related to thermostability based on scikit-learn, there are a total of 10297 mutations and 169 sequences in the data. In order to test the performance of CASPE, we selected 10 sites for each sequence by Grad-CAM or random screening, and the amino acid microenvironments were obtained. Finally, the amino acid microenvironment was regarded as input to obtain the probability of 20 amino acids’ occurrence at this site. There are 1674 sites and 1690 sites for Grad-CAM and random screening, respectively. When we selected sites by Grad-CAM, some sites’ amino acid microenvironments may be discarded due to being too large or too small to adapt to the model, thereby obtaining a total of 1674 sites. Based on the data selected above, we performed PCA analysis and obtain the top 5 main components.

### Interpretability analysis of APCNet

To further explore the importance of the three microenvironments, we selected a fixed database and generated data containing all three microenvironments, data for microenvironments without atoms, data for microenvironments without charges, and data for microenvironments without SASA. We trained the model to explore the impact of various microenvironments on model training. Moreover, three interpretability techniques, including SHAP^82^ (SHapley Additive exPlanations), Captum^83^, and Permutation Importance^84^ were adopted to fully analyze the logic of black-box models. Specifically, we used the validation and test sets from model training to interpret the model. SHAP calculates the contribution of features based on Shapley values. We adopted the shap.GradientExplainer in Python to calculate SHAP values for all samples. Captum is Facebook’s official interpretability tool for models based on the PyTorch framework. We adopted the captum.attr in Python to calculate Captum values for all samples. Permutation Importance measures the ranking importance of all features in a model by calculating the difference in weights before and after shuffling features. Because the order of microenvironments has no influence on model training, we shuffled the features by adding a random number between 0 and 1 to their original values.

## Supporting information

https://drive.google.com/file/d/1yqwkn4ss5dsXfbrrVM4BBC9nM6JZDZFD/view?usp=drive_link

## Acknowledgments

We sincerely thank Prof. Lijiao Tian for revising the article, Prof. Huimin Zhao and Singh Nilmani for providing phytase plasmid map.

## Author contributions

Conceptualization: Q. Deng, C. Wang, M. Jin, X. Li, H. Cui; Methodology: Q. Deng, J. Qiao, C. Wang, R. Zhai, X. Li, M. Jin; Investigation: Q. Deng, J. Qiao, C. Wang, X. Ni, Y. Chang, N. Zhao; Visualization: Q. Deng, C. Wang; Funding acquisition: M. Jin, X. Li; Project administration: Q. Deng, X. Li, M. Jin; Supervision: X. Li, M. Jin; Writing – original draft: Q. Deng, C. Wang; Writing – review & editing: Q. Deng, H. Cui, M. Jin, R. Zhai.

## Code availability

All code in this work can be obtained for free. Code for CASPE can be freely accessible at GitHub (https://github.com/Yusei-Star/CASPE).

## Supplementary Materials

Supplementary Text

Figs. S1 to S42

Tables S1 to S2

References (1-7)

