## Supplementary material for "Discriminative Site-Directed Protein Engineering via Lightweight CASPE Platform": https://drive.google.com/file/d/1yqwkn4ss5dsXfbrrVM4BBC9nM6JZDZFD/view?usp=drive_link: supporting_information_0423.pdf

**A Critical Sites-Driven and Light-weighted Protein Engineering Platform**

[#] These authors contributed equally.

**This PDF file includes:**

Supplementary Text
Figs. S1 to S42
Tables S1 to S3
References (1-5)

Table of Contents

**Fig. S 1 | The number of protein sequences in different temperature ranges above 60℃ in the learn2therm database.** ......9
**Fig. S 2 | Overview of CASPE and its applications.** (a) The structure of protein classification model, and the model is generated by transfer learning, ESM2 is connected by MLP. (b) The process of obtaining the key amino acid sites based on CAS. (c) The process of predicting the optimal amino acid by APCNet. (d) The process of model validation, there are two steps including comparing with other models and wet-lab experimental tests (This figure was generated by biorender, https://app.biorender.com). ......10
**Fig. S 16 | The conservation analysis of all mutation sites selected by CAS of BG,**

**EG and CBHI based on the model related to thermostability.** The complete list of mutations for **BG** is D136A, G137G, G138G, W139Y, A140A, S141S, T144T, A145A, H146F, D164D, A165A, V166V, L196L, A197A, A198A, N227A, A228A, H229F, A239A, D240N, E267E, M268M, M269M, E270E, A271A, L272L, N297A, Y298Y, Y299Y, T300T, P301P, V304V, A305A, D306A, Q319A, A320A, P321A, Y335Y, A336A, P337P, A338A, L339L, E351E, L352L, P353P; **EG:** N42D, T43T, T44E, A45N, I90I, T91I, G92S, S149N, V150L, V151V, V152V, A153S, L154L, Q156Q, A157S, A158A, A234T, T235T, T236Q, W237F, A251G, G252G, G253G, A254N; **CBHI:** C19S, S20S, S57S, S58S, G75G, A76A, A77A, Y78F, A79A, S80T, D114S, T115T, T116T, Q186Q, A187A, N188T, T201T, G202G, I203T, G204G, G205G, H206N, G207G, S208S, Q312S, Q313L, P314T, C331G, T332G, A333S, E334Q, A355A, T356T, S357G, S400T, S401S, G402G........................................................24
**Fig. S 21 | The conservation analysis of all mutation sites selected by CAS of BG based on CASPEA.** The complete list of mutations for acidic stability is G13G, D14G, F15F, I90L, I91I, P92P, I118I, K119K, T120Q, S235S, D236G, S237S, A239G, D240N, L241Q, A266G, E267Q, M268M, M269M, E270E, A305A, D306D, D307D, Y350Y, E351D, L352L; **alkaline:** G13G, D14G, F15F, A42Q, F43Y, C44G, D68E, D69D, L70L, I90I, I91I, P92P, I118I, K119K, T120T, S235S, A239S, D240D, L241E, F260F, K261G, G262G, A266G, E267E, M268M, M269M, E270E, A305A, D306D, D307D, Y350Y, E351D, L352L, E372E, V373V, P377P, R378R, L379Q. ...........................29
**Fig. S 26 | Specific activity of mutants relative to wild-type, and the mutants were**

**Fig. S 41 | Count the ranking of each feature among the 20 kinds of amino acids (Captum).** (a) for thermostability, (b) for acid stability and (c) for alkaline stability. 49

### Supplementary Text

#### Supplementary Text 1. CASPE evaluation *in silico* and interpretability analysis of APCNet

For PC1 (Fig. 5e and Fig. S36), based on the comprehensive analysis of amino acid hydrophobicity, charge, the rigidity of protein structure and other characteristics, the amino acids play a positive role in PC1 focus on "intrinsic stability", such as the core amino acids of hydrophobicity (I, V, L and M), which are helpful to forming the internal core of the protein. Moreover, R and K are positively charged, which are beneficial to forming ionic bonds and stabilizing the protein structure. The amino acids play a negative role in PC1 focus on "extrinsic adaptability", and the flexible side chains of G and A enable changes in protein conformation, the same as polar/charged amino acids (D, E, H, N) [5]. The amino acids play a positive role in PC2 focus on "catalytic function" (acidic/thiol group) and "structural regulation" (proline and hydrophobic core). The amino acids play a negative role in PC2 focus on "dynamic adaptability" (flexible surface, aromatic ring signal) and "ion regulation" (basic amino acids). Based on PC1 and PC2, the thermostability of proteins focuses on choosing hydrophobic amino acids that are conducive to the formation of stable intrinsic properties, rather than amino acids with high flexibility.

#### Supplementary Text 2. Limitations of using MLM for zero-shot mutation effect prediction

***MLM prediction as conditional probability***

Methods for predicting mutational effects of masked language models (MLMs, such as ESM-2) is to predict hidden (masked) amino acids based on the surrounding sequence context. Mathematically, for a protein sequence $X$ of length $L$, we denote the amino acid at position $i$ as $x_i$, and the remaining sequence context (excluding position $i$) as $X_{-i}$. Strictly, the predicted output of MLM at position $i$ is a conditional probability:

$$P(x_i|X_{-i})$$

According to the basic definition of joint probability, the probability $P(X)$ of a full-length sequence $X$ appearing in nature can be expressed as the joint probability of the target site $X_i$ and its context $X_{-i}$:

$$P(X) = P(x_i|X_{-i}) \cdot P(X_{-i})$$

$$P(x_i|X_{-i}) = \frac{P(X)}{P(X_{-i})}$$

Taking the logarithm of both sides of the equation, we can get the output of MLM:

$$logP(x_i|X_{-i}) = logP(X) - logP(X_{-i})$$

The raw probability score $logP(x_i|X_{-i})$ predicted by MLM is adjusted by $logP(X_{-i})$, which represents the probability that the sequence context exists in nature. Mathematically, comparing the raw MLM output probabilities of different sites (e.g., site $i$ vs. site $j$) needs further investigation because the context probabilities $P(X_{-i})$ and $P\left(X_{-j}\right)$ on which they are based are different.

#### *Mathematical derivation of* $\Delta LL_i$

To assess the mutational effect at site $i$ without contextual bias, the zero-shot method in the literature[1] defines the mutational effect as the log probability difference ($\Delta LL_i$ ) between the mutated residue $x_i^{mut}$ and the wild-type residue $x_i^{wt}$, given the same wild-type context $X_{-i}^{wt}$:

$$\Delta LL_i = logP(x_i^{mut}|X_{-i}^{wt}) - logP(x_i^{wt}|X_{-i}^{wt})$$

Substituting the conditional probability relation derived in "***MLM prediction as conditional probability***" into this equation, we obtain:

$$\Delta LL_i = \left(logP\left(X^{mut\_i}\right) - logP(X_{-i}^{wt})\right) - \left(logP(X^{wt}) - logP(X_{-i}^{wt})\right)$$

Where, $X^{mut\_i}$ is the sequence after mutating into a new residue at position $i$, and $X^{wt}$ is the wild-type sequence.

By eliminating the common context term $logP(X_{-i}^{wt})$ to simplify the equation, we obtain:

$$\Delta LL_i = logP\left(X^{mut\_i}\right) - logP(X^{wt})$$

This simplified equation demonstrates that the bias introduced by the specific structural context $X_{-i}^{wt}$ can be effectively offset by calculating the log-likelihood ratio relative to wild-type residues. The resulting $\Delta LL_i$ is mathematically rigorously equivalent to the difference in the global joint probability between the mutant and wild-type sequences in the natural evolution.

#### *Derivation and Simplification of* $\Delta LL_i - \Delta LL_j$

When we need to compare the mutational effects occurring at two different sites $i$ and $j$ of the same sequence $X^{wt}$, we evaluate the difference between their respective $\Delta LL$ scores:

$$\Delta LL_i - \Delta LL_j = \left(logP\left(X^{mut\_i}\right) - logP(X^{wt})\right) - \left(logP\left(X^{mut\_j}\right) - logP(X^{wt})\right)$$

Expanding this equation, the joint probability term for wild types is perfectly canceled out:

$$\Delta LL_i - \Delta LL_j = logP\left(X^{mut\_i}\right) - logP(X^{wt}) - logP\left(X^{mut\_j}\right) + logP(X^{wt})$$

$$\Delta LL_i - \Delta LL_j = logP\left(X^{mut\_i}\right) - logP\left(X^{mut\_j}\right)$$

Using the quotient rule of logarithms, the final simplified form is:

$$\Delta LL_i - \Delta LL_j = log\left(\frac{P\left(X^{mut\_i}\right)}{P\left(X^{mut\_j}\right)}\right)$$

The final results demonstrate that using $\Delta LL$ scores to assess whether a mutation at site $i$ is superior to a mutation at site $j$ is mathematically rigorously equivalent to comparing the ratio of their full-sequence joint probabilities $P\left(X^{mut\_i}\right)$ to $P\left(X^{mut\_j}\right)$ , this means that the larger $P(X^{mut})$is, the higher the $\Delta LL$ score.

While mathematical simplification establishes a unified benchmark for cross-site comparisons, Mathematical simplification establishes a unified benchmark for cross-site comparisons, it reveals a limitation of unsupervised zero-shot prediction in protein engineering. $\Delta LL$ rankings depends on the joint distribution *P(X)* learned from unlabeled natural protein databases (such as UniRef50).
The *P(X)* of sequence statistical significance only characterizes the stability of the sequence in maintaining protein function and structure during natural evolutionary selection[2,3]. Therefore, it cannot bridge the gap between natural evolution and industrial applications[4]. Overall, $\Delta LL_i - \Delta LL_j$ is an indicator used to maintain protein stability. Without combining supervised learning labeled data or biophysical structural constraints, it has limitations in selecting protein mutation sites for industrial applications.

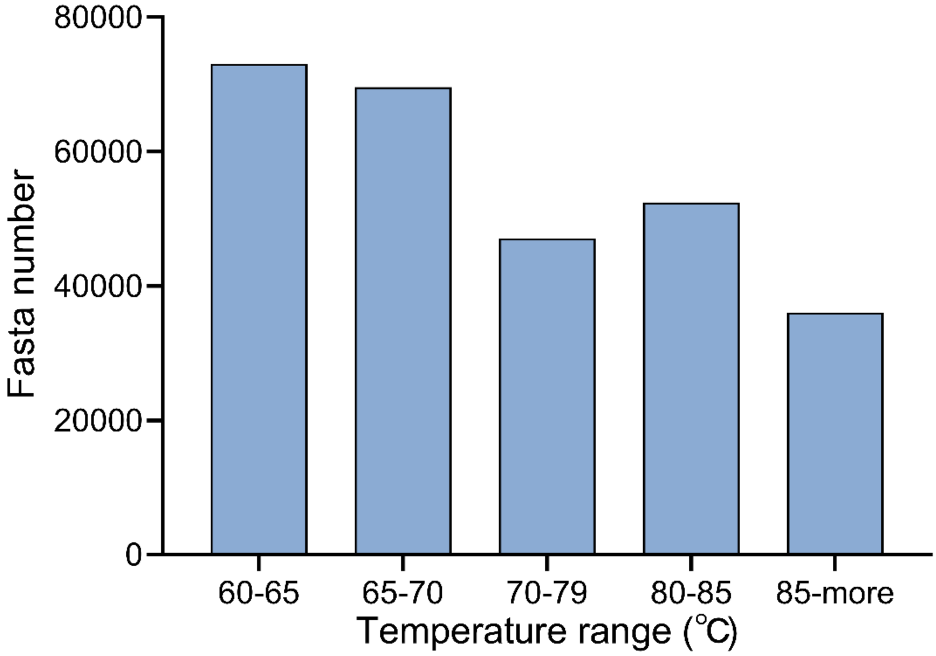

**Fig. S 1 | The number of protein sequences in different temperature ranges above 60°C in the learn2therm database.**

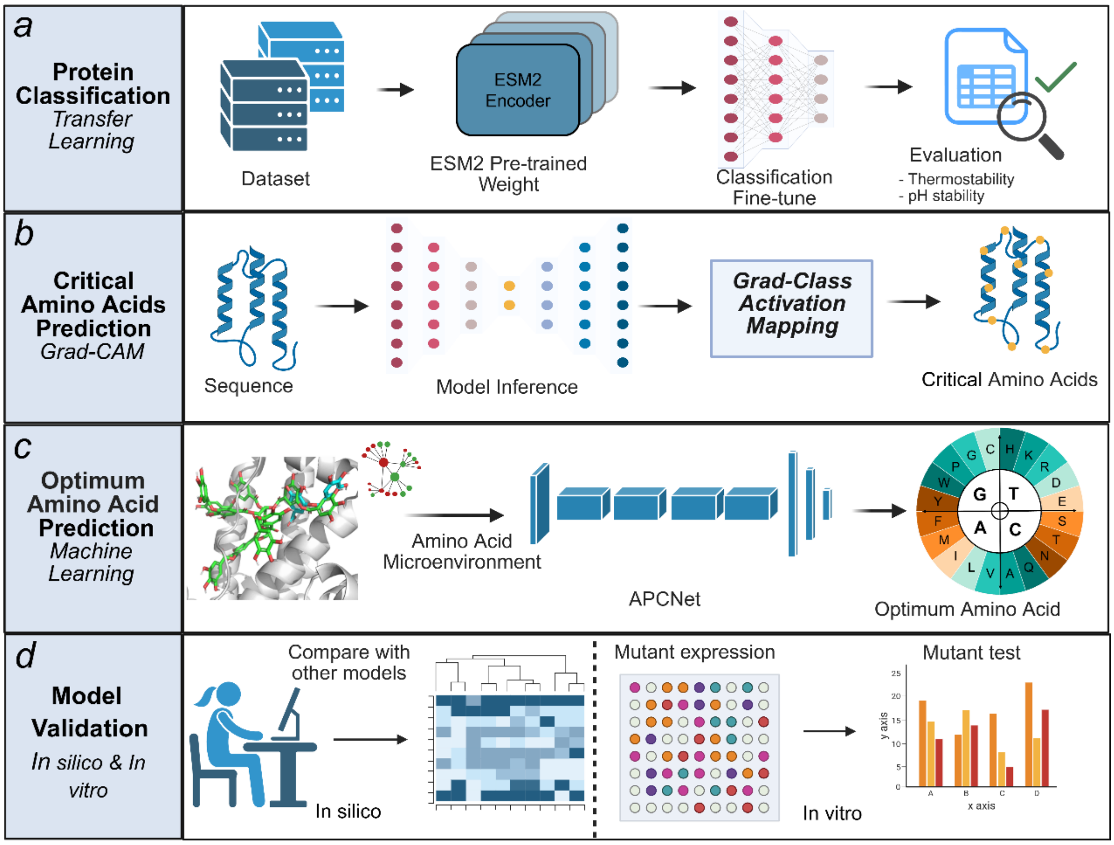

**Fig. S 2 | Overview of CASPE and its applications.** (a) The structure of protein classification model, and the model is generated by transfer learning, ESM2 is connected by MLP. (b) The process of obtaining the key amino acid sites based on CAS. (c) The process of predicting the optimal amino acid by APCNet. (d) The process of model validation, there are two steps including comparing with other models and wet-lab experimental tests (This figure was generated by biorender, https://app.biorender.com).

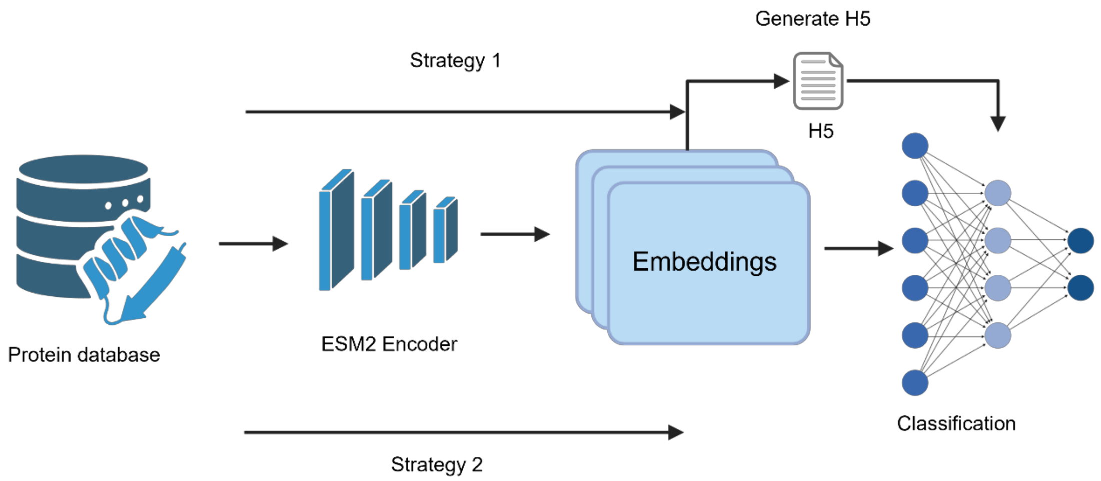

**Fig. S 3 | Two strategies based on ESM2 in this experiment.** Strategy 1: the embeddings files are generated by ESM2 and then put them into a CNN classification model for training; Strategy 2: attach the classification layer directly after ESM2 generates high-dimensional vectors to skip the process of generating the embedding file.

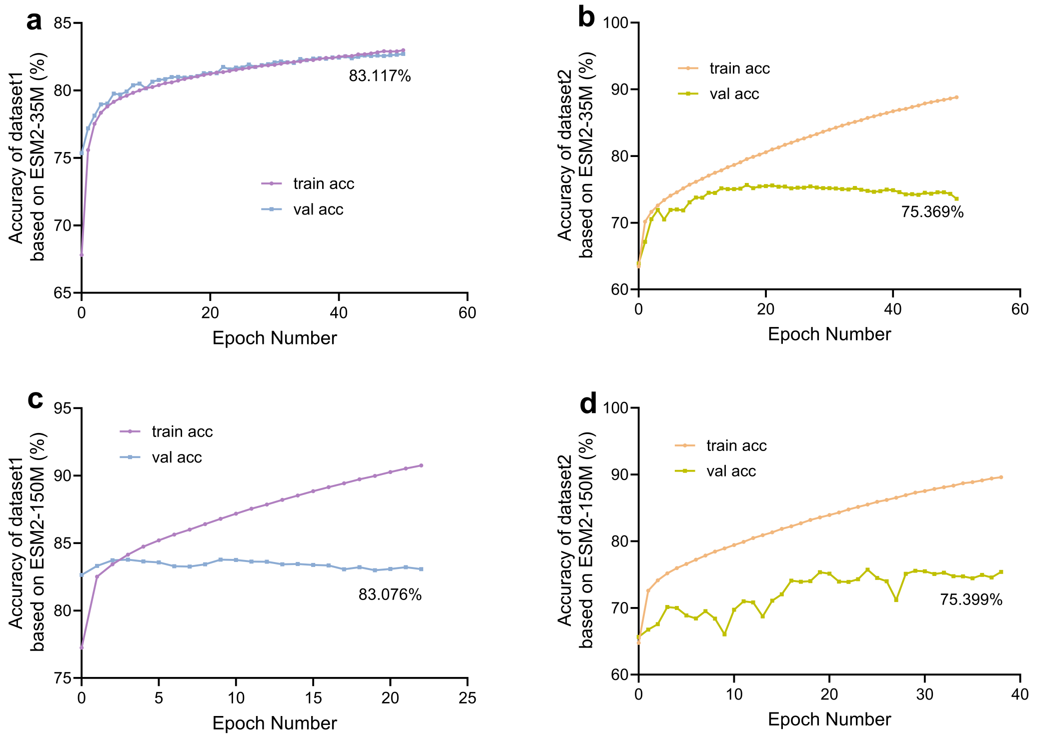

**Fig. S 4 | Training results based on esm2_t12_35M and esm2_t30_150M.**

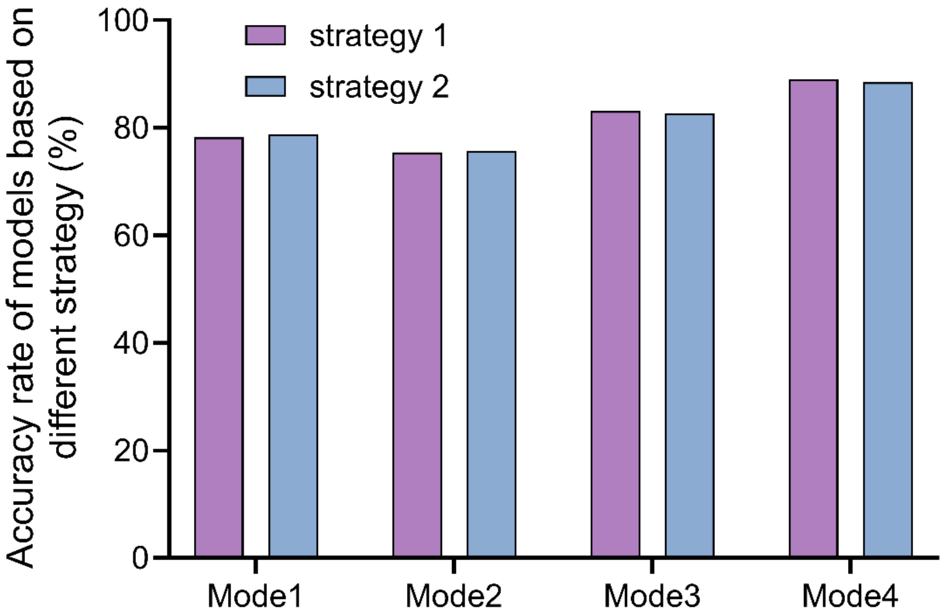

**Fig. S 5 | The results of two strategies based on ESM2 in this experiment.**

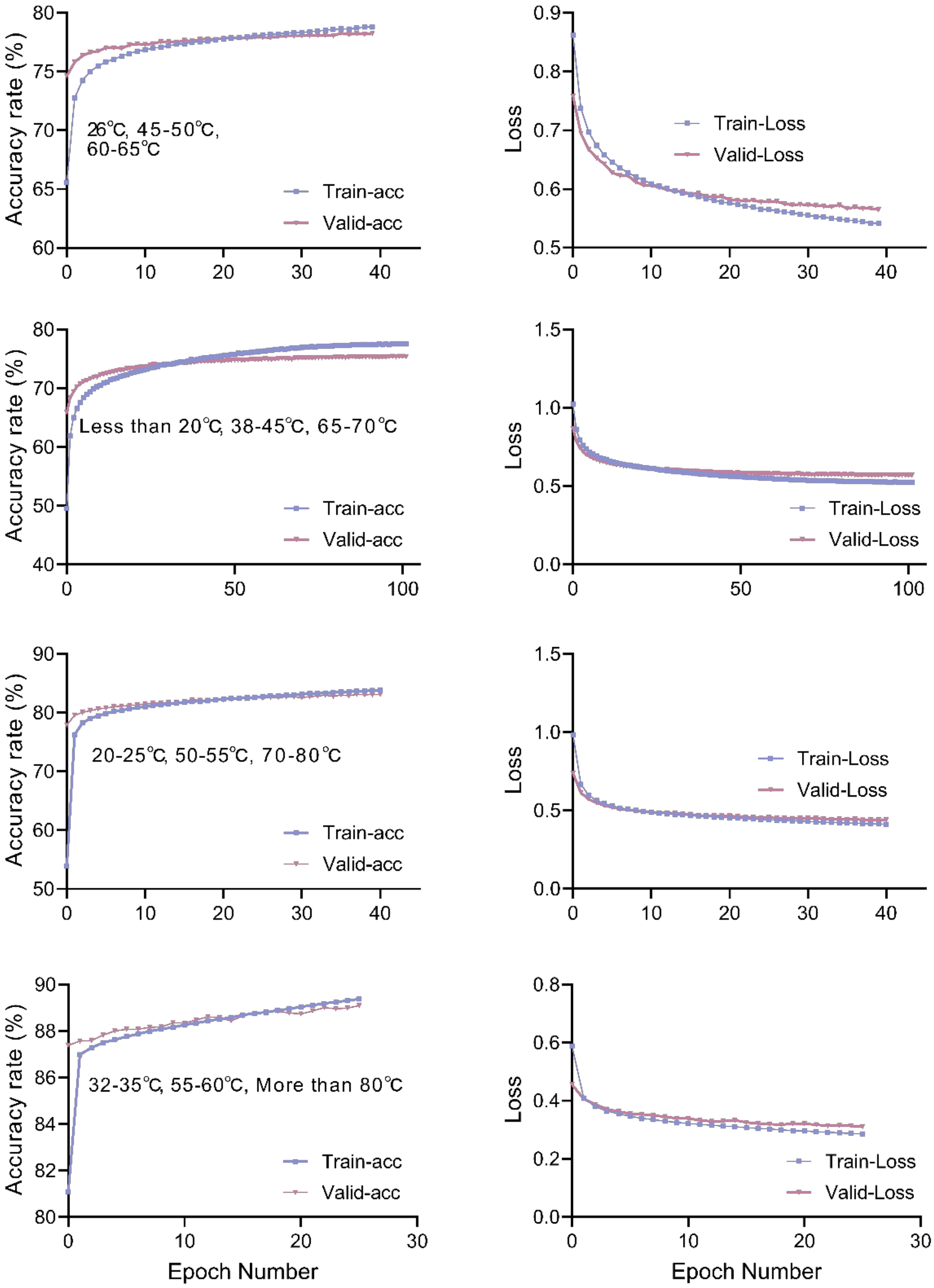

**Fig. S 6 | The results of strategy 1 based on ESM2 in this experiment.**

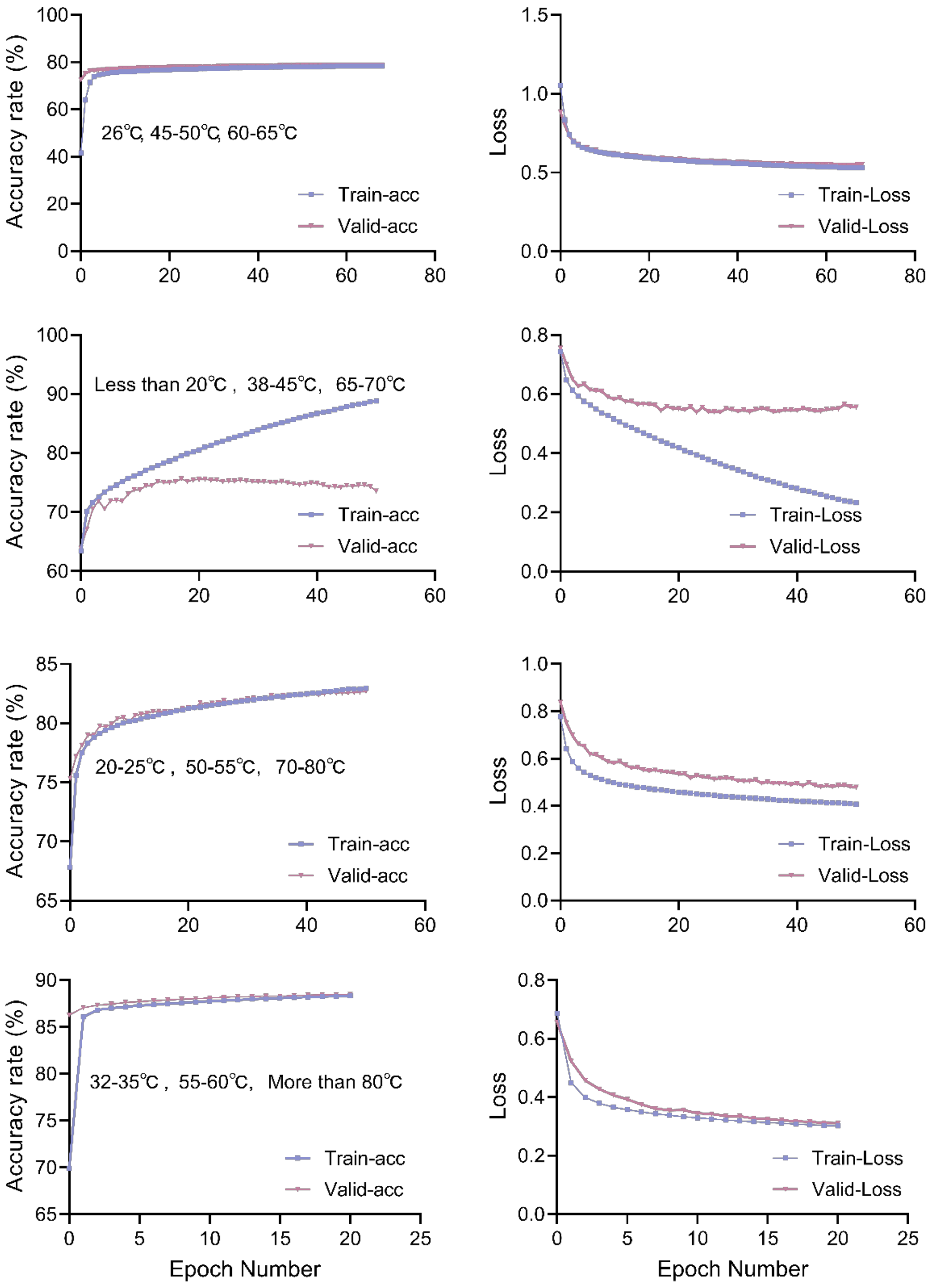

**Fig. S 7 | The results of strategy 2 based on ESM2 in this experiment.**

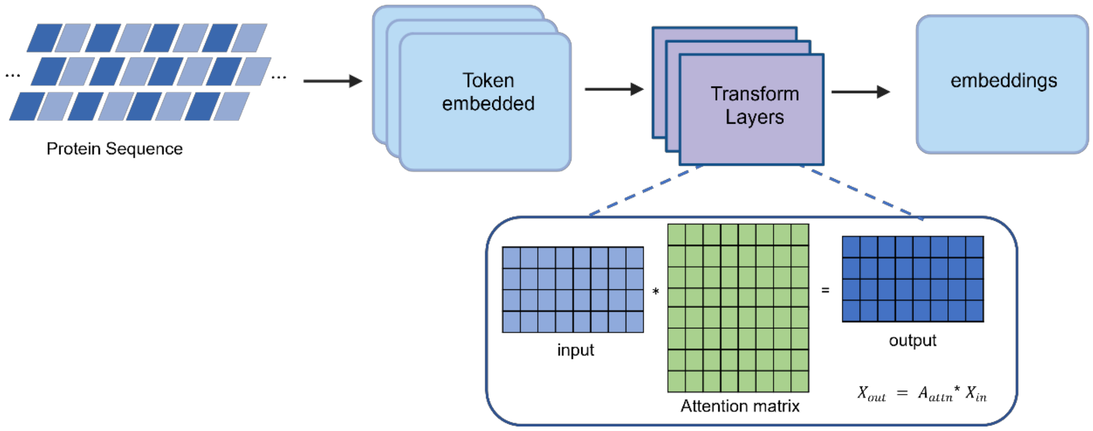

**Fig. S 8 | ESM2 data processing.**

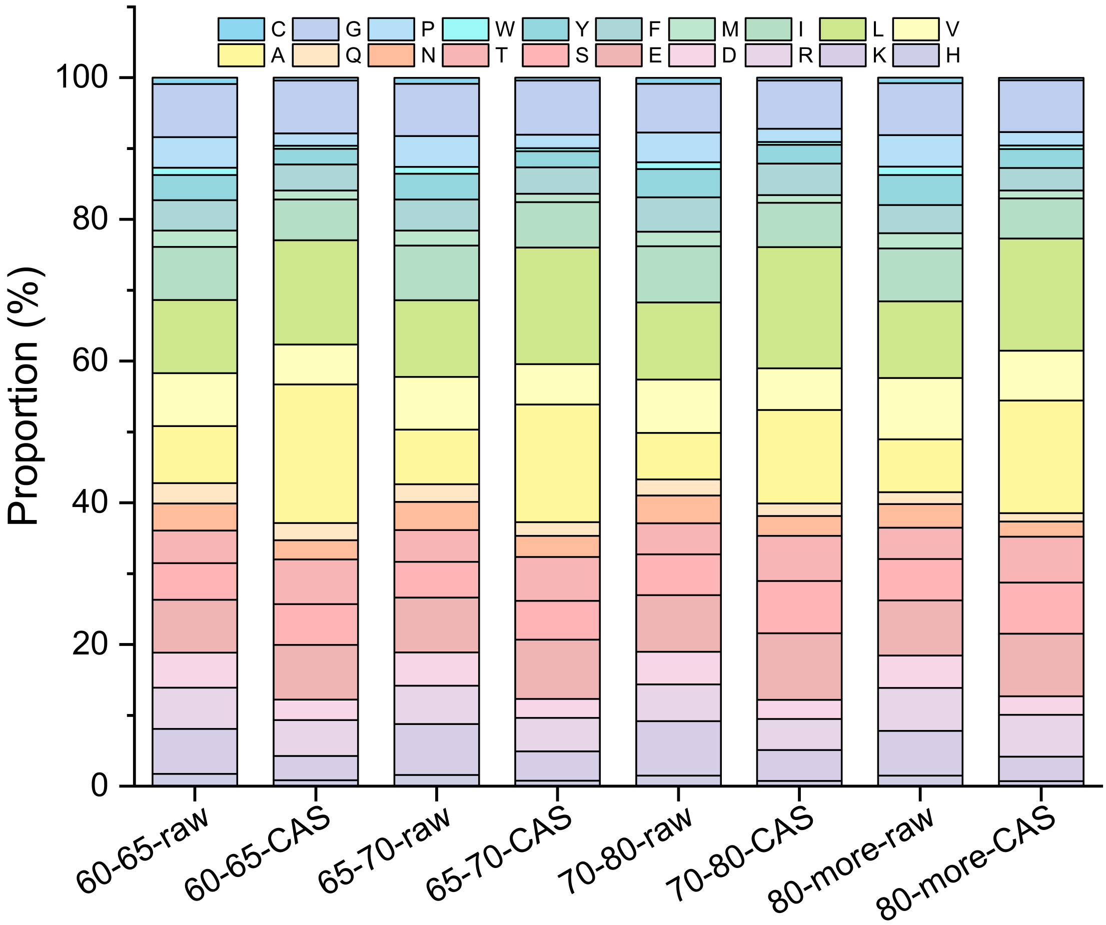

**Fig. S 9 | The proportion of standard amino acids in protein sequences related to thermostability before and after CAS.**

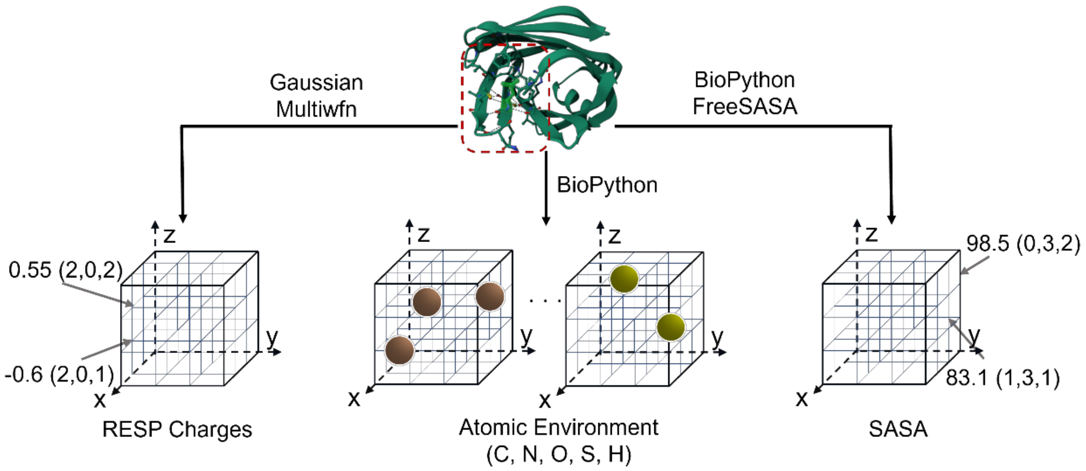

**Fig. S 10 | Schematic diagram of microenvironment data sampling process.**

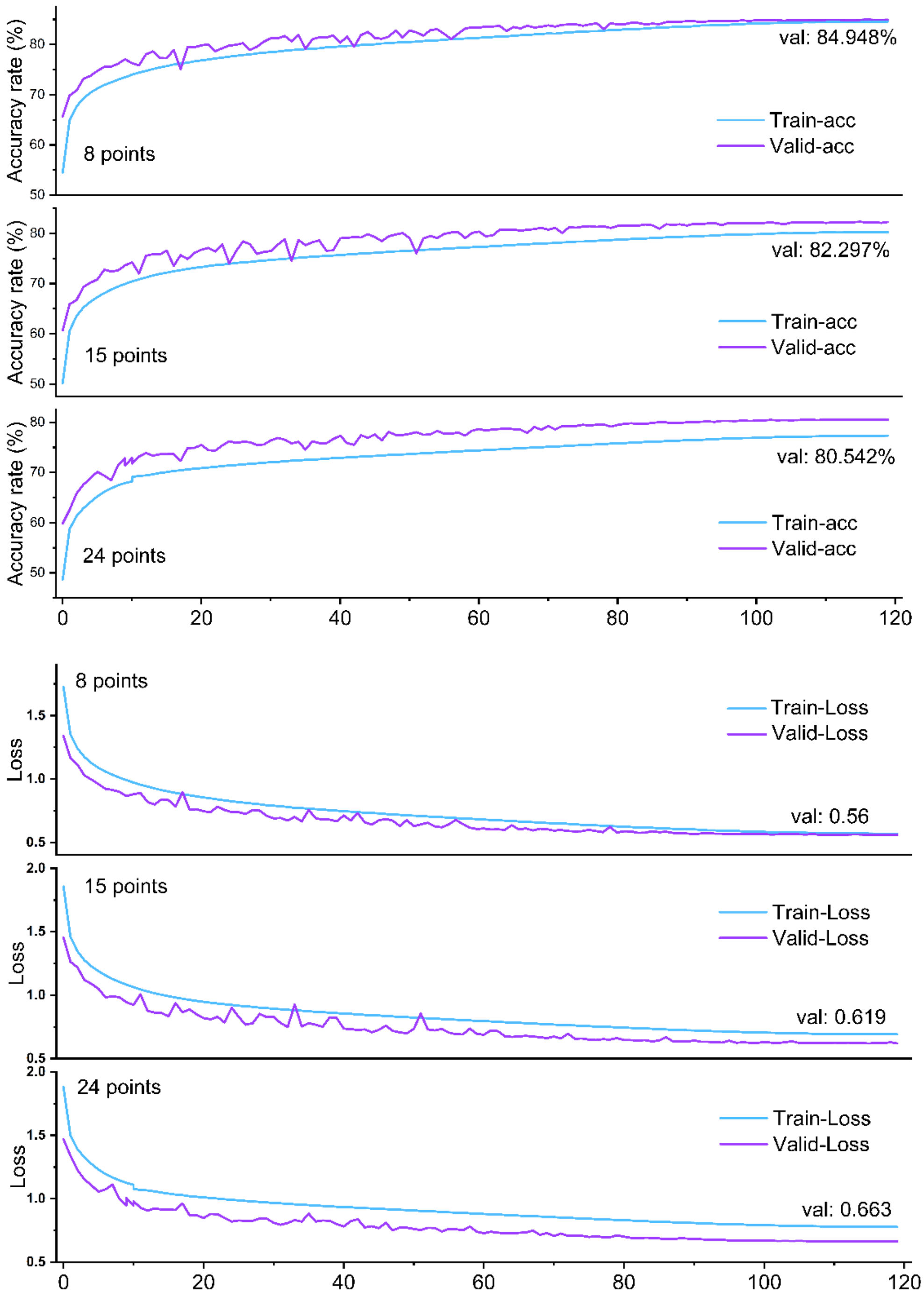

**Fig. S 11 | Training results of APCNet based on protein sequences related to thermostability.** 8 points: we selected 8 critical amino acids for each sequence which are relevant to thermostability selected by CAS; 15 points: 5 critical amino acids plus 10 adjacent residues before/after each critical site for each sequence; 24 points: 8 critical amino acids plus 16 adjacent residues before/after each critical site for each sequence.

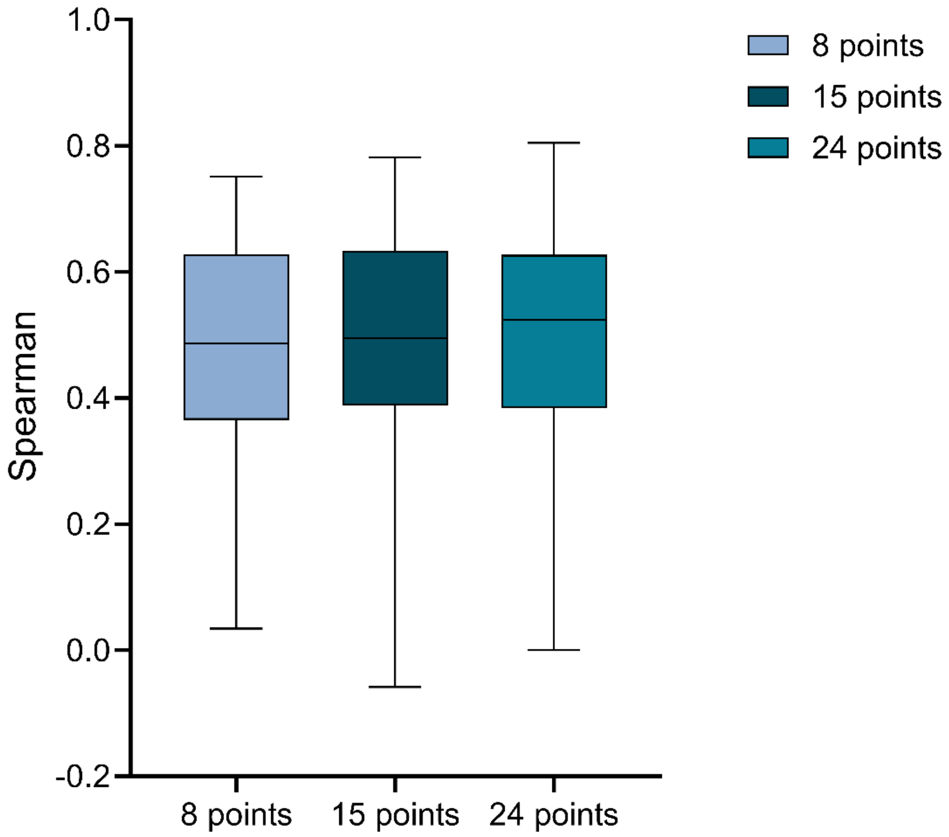

**Fig. S 12 | The spearman of different APCNet model based on CDNA.** 8 points: we selected 8 critical amino acids for each sequence which are relevant to thermostability selected by CAS; 15 points: 5 critical amino acids plus 10 adjacent residues before/after each critical site for each sequence; 24 points: 8 critical amino acids plus 16 adjacent residues before/after each critical site for each sequence.

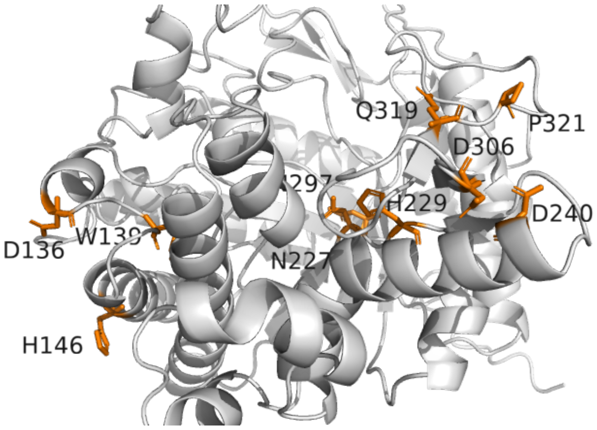

**BG**

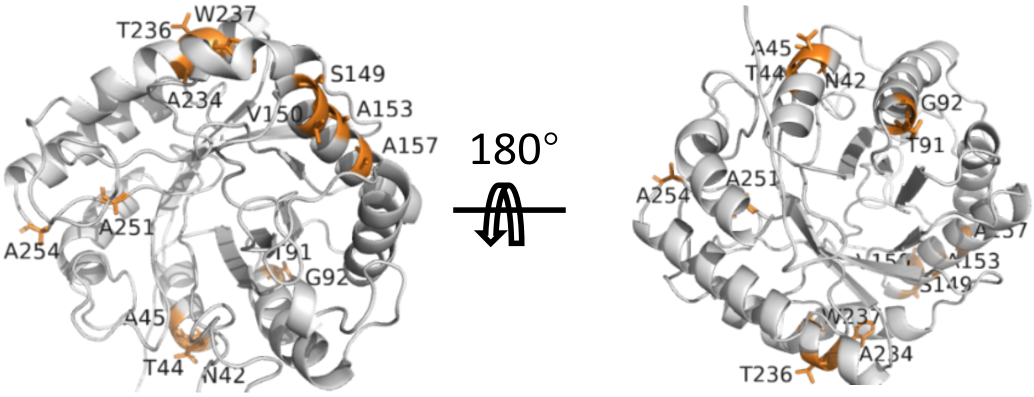

**EG**

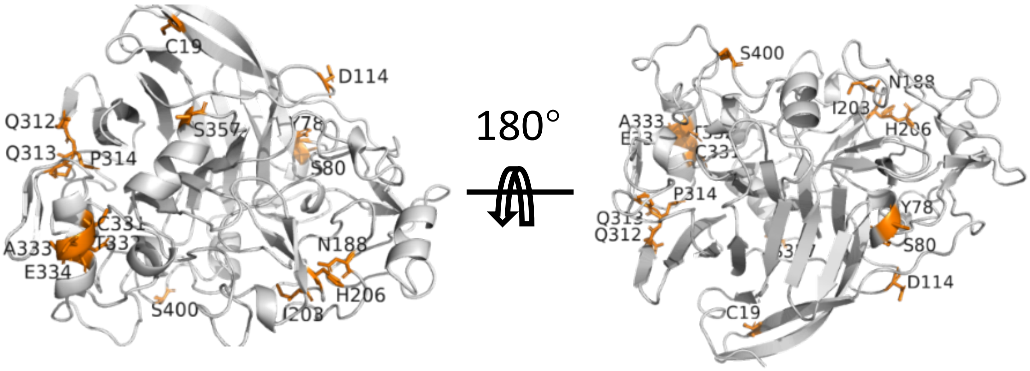

**CBHI**

**Fig. S 13 | Mutation sites selected by CASPET of BG, EG and CBHI.**

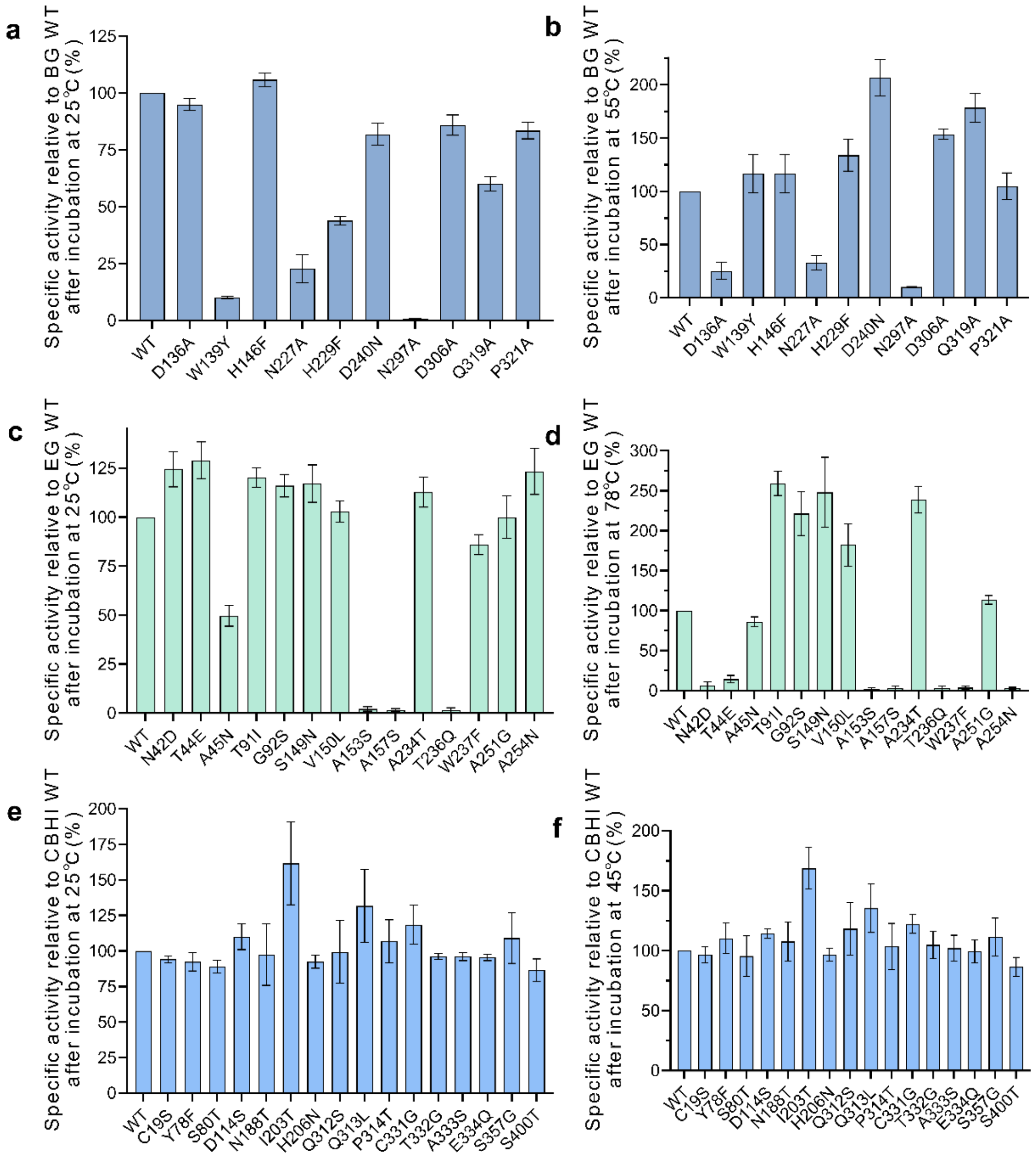

**Fig. S 14 | Specific activity relative to wild-type after incubate at 25 °C or high temperature** (BG: 55 °C, EG: 78 °C, CBHI: 45 °C). BG: (a, b); EG: (c, d); CBHI: (e, f).

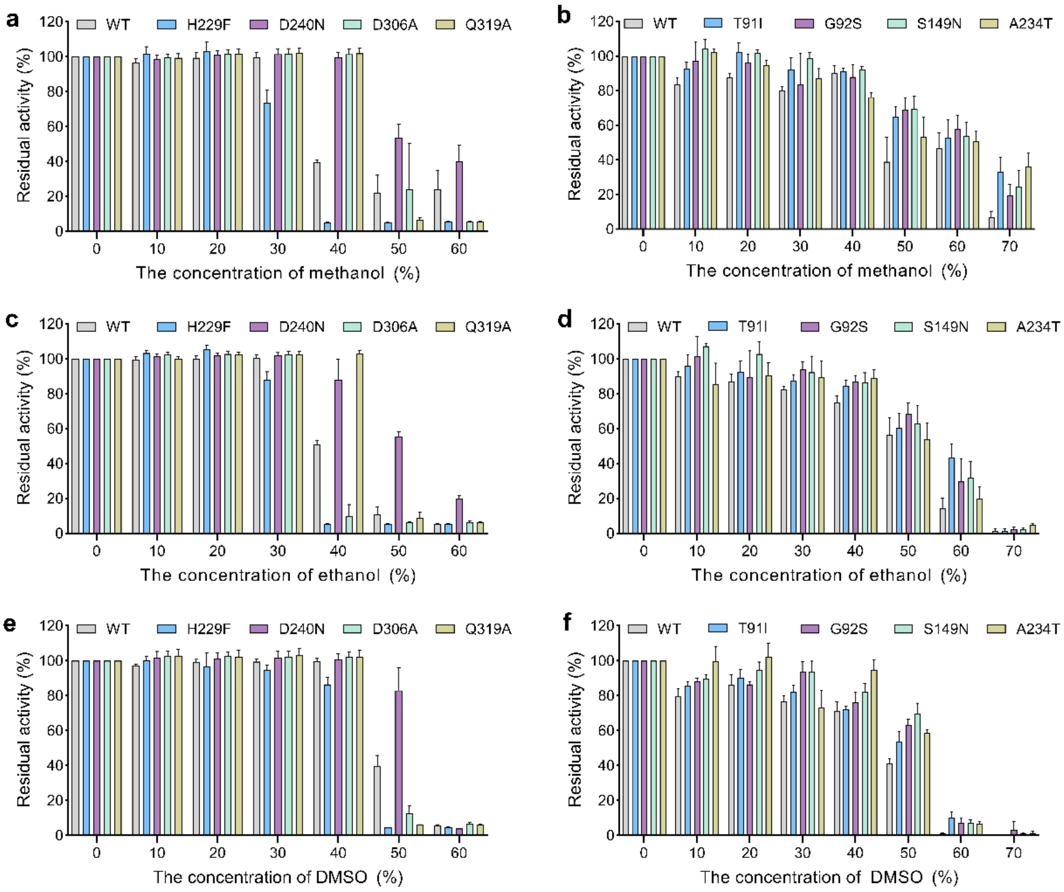

**Fig. S 15 | Specific activity relative to wild-type after incubate at methanol, ethanol or DMSO.** (a, c, e) for BG and (b, d, f) for EG. (a, b) The relative activity of enzymes under different concentrations of methanol. (c, d) The relative activity of enzymes under different concentrations of ethanol. (e, f) The relative activity of enzymes under different concentrations of DMSO.

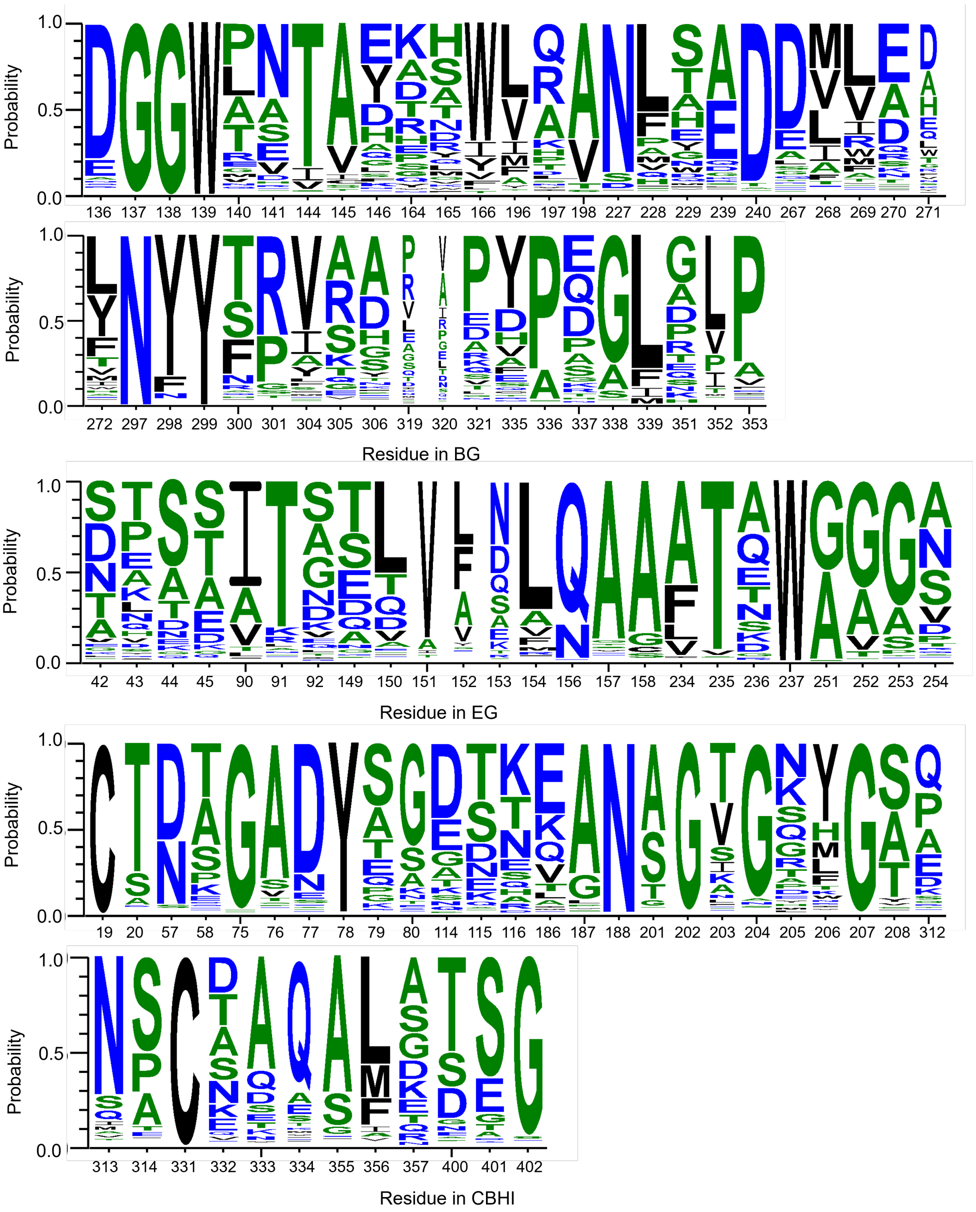

**Fig. S 16 | The conservation analysis of all mutation sites selected by CAS of BG, EG and CBHI based on the model related to thermostability.** The complete list of mutations for **BG** is D136A, G137G, G138G, W139Y, A140A, S141S, T144T, A145A, H146F, D164D, A165A, V166V, L196L, A197A, A198A, N227A, A228A, H229F, A239A, D240N, E267E, M268M, M269M, E270E, A271A, L272L, N297A, Y298Y, Y299Y, T300T, P301P, V304V, A305A, D306A, Q319A, A320A, P321A, Y335Y, A336A, P337P, A338A, L339L, E351E, L352L, P353P; **EG:** N42D, T43T, T44E, A45N, I90I, T91I, G92S, S149N, V150L, V151V, V152V, A153S, L154L, Q156Q, A157S, A158A, A234T, T235T, T236Q, W237F, A251G, G252G, G253G, A254N; **CBHI:** C19S, S20S, S57S, S58S, G75G, A76A, A77A, Y78F, A79A, S80T, D114S, T115T, T116T, Q186Q, A187A, N188T, T201T, G202G, I203T, G204G, G205G, H206N, G207G, S208S, Q312S, Q313L, P314T, C331G, T332G, A333S, E334Q, A355A, T356T, S357G, S400T, S401S, G402G.

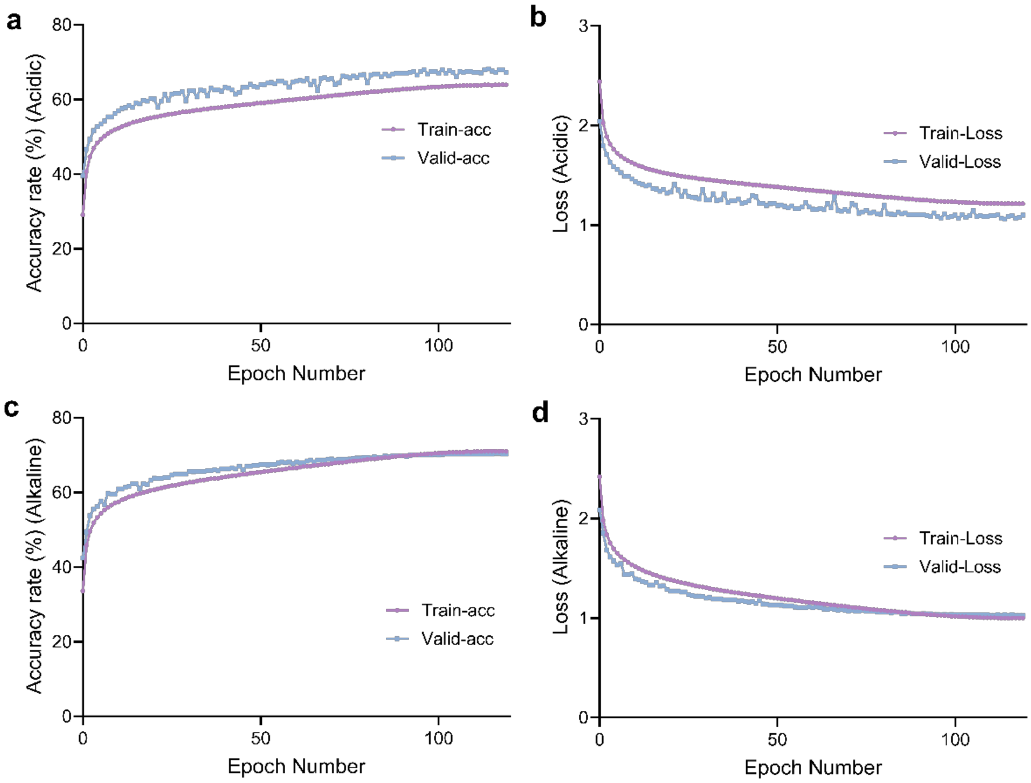

**Fig. S 17 | Train results of APCNet based on protein sequences related to pH stability.**

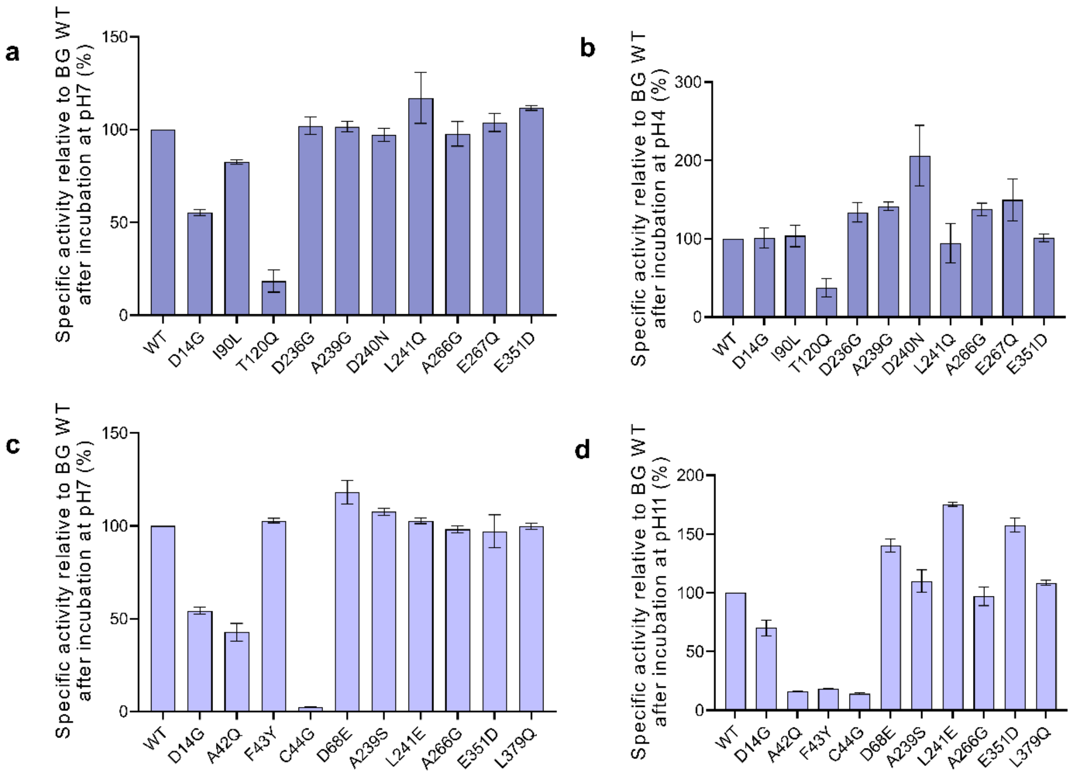

**Fig. S 18 | Specific activity relative to wild-type after incubate at pH 4 and pH 11.** (a, b) The relative activity of enzymes under neutral or acidic conditions. (c, d) The relative activity of enzymes under neutral or alkaline conditions.

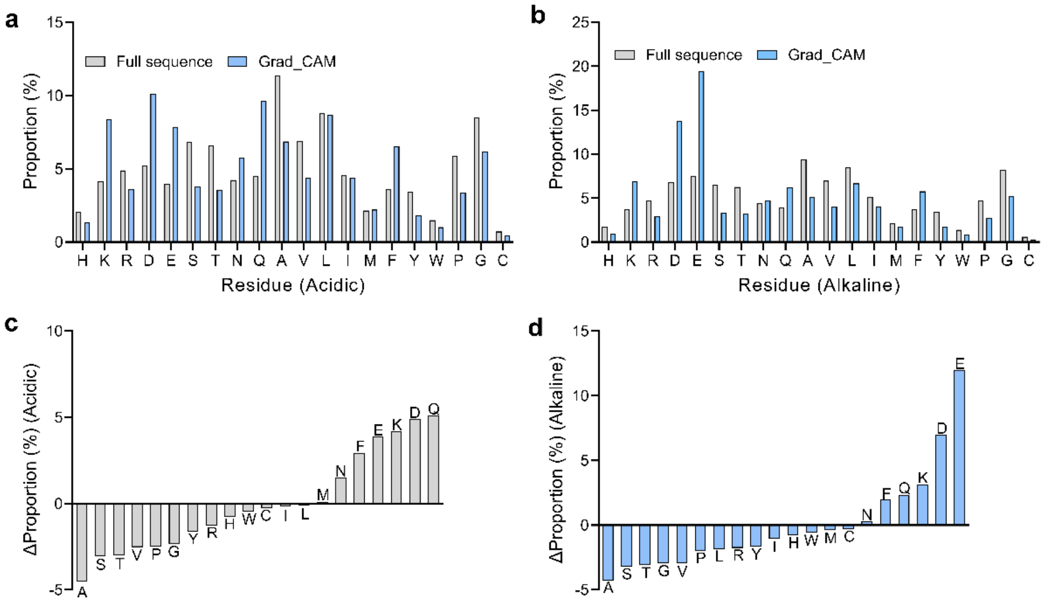

**Fig. S 19 | Amino acid proportions before and after the CAS.** (a, b) The proportion of standard amino acids in protein sequences related to pH stability before and after CAS. (c, d) The Δ proportion of standard amino acids in protein sequences related to pH stability before and after CAS (a, c) for acid stability and (b, d) for alkaline stability.

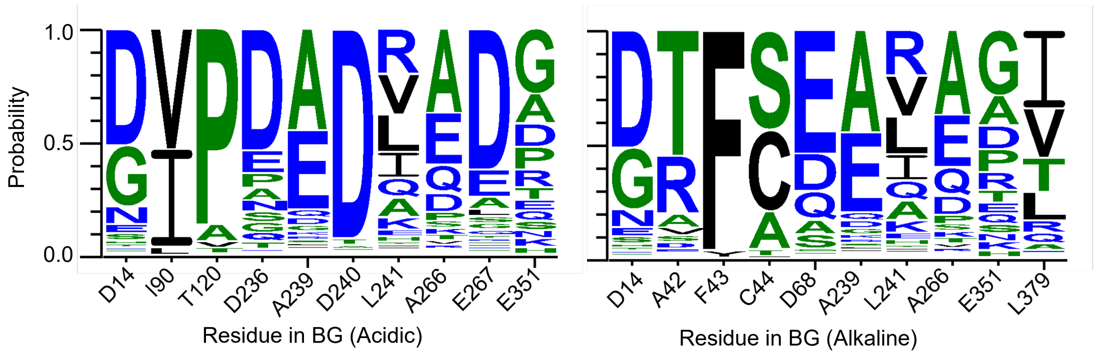

**Fig. S 20 | Conservation analysis of residues of BG based on the model related to pH stability (obtained from WebLogo3 analysis).**

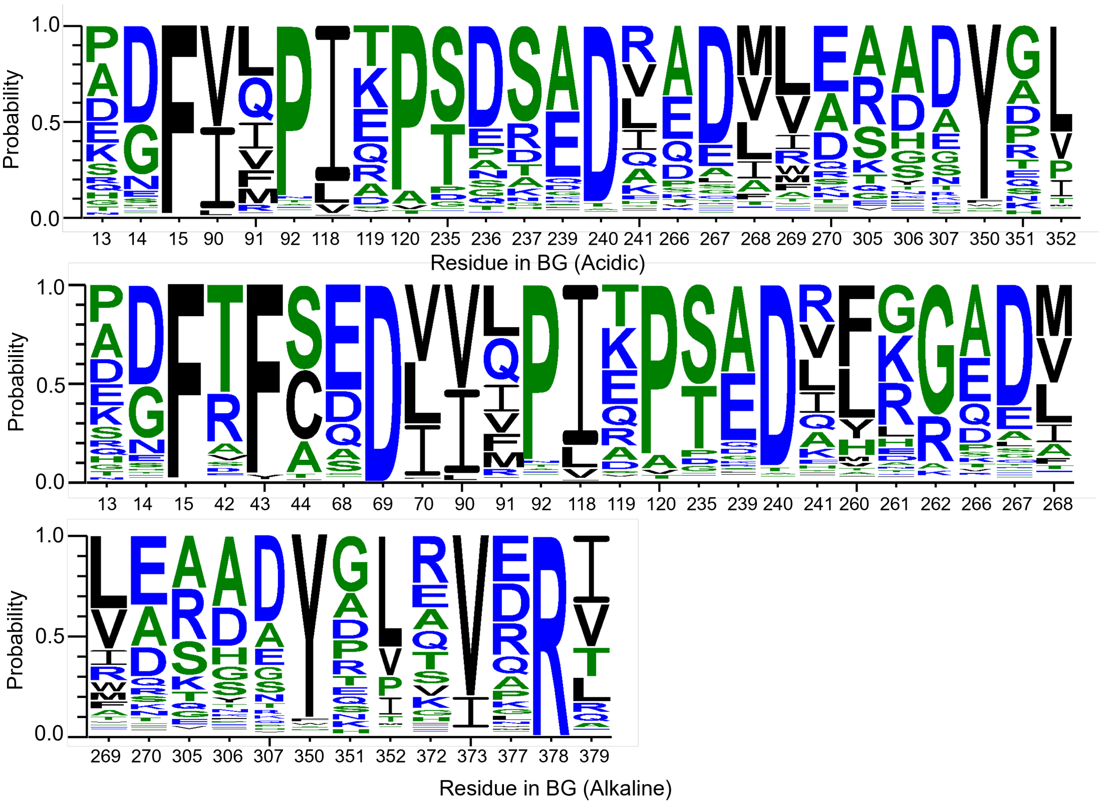

**Fig. S 21 | The conservation analysis of all mutation sites selected by CAS of BG based on CASPEA.** The complete list of mutations for acidic stability is G13G, D14G, F15F, I90L, I91I, P92P, I118I, K119K, T120Q, S235S, D236G, S237S, A239G, D240N, L241Q, A266G, E267Q, M268M, M269M, E270E, A305A, D306D, D307D, Y350Y, E351D, L352L; **alkaline:** G13G, D14G, F15F, A42Q, F43Y, C44G, D68E, D69D, L70L, I90I, I91I, P92P, I118I, K119K, T120T, S235S, A239S, D240D, L241E, F260F, K261G, G262G, A266G, E267E, M268M, M269M, E270E, A305A, D306D, D307D, Y350Y, E351D, L352L, E372E, V373V, P377P, R378R, L379Q.

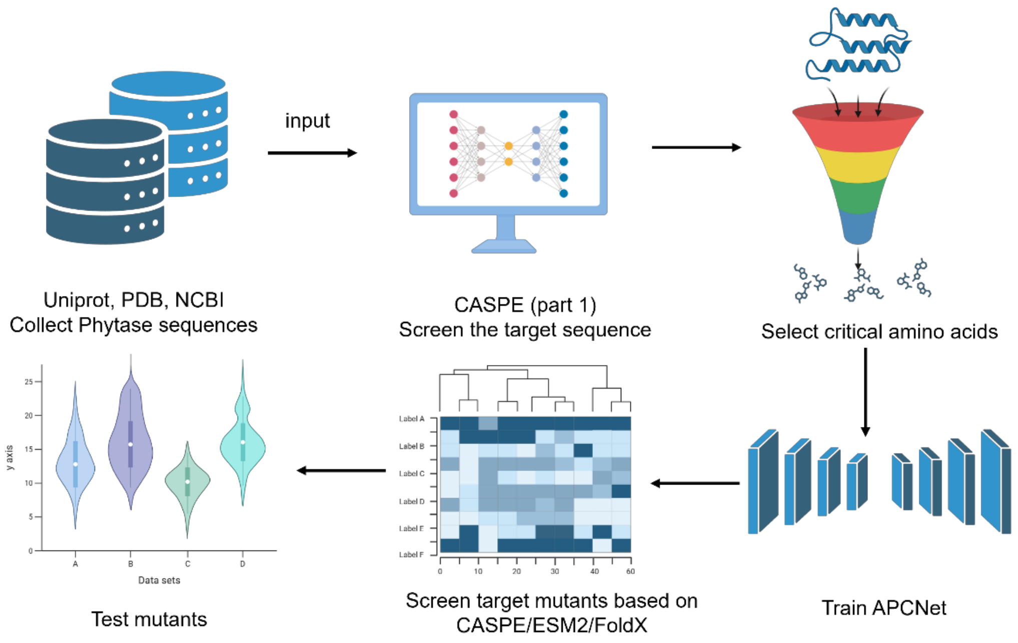

**Fig. S 22 | The framework of model verification based on phytase sequences** (This figure was generated by biorender, https://app.biorender.com/).

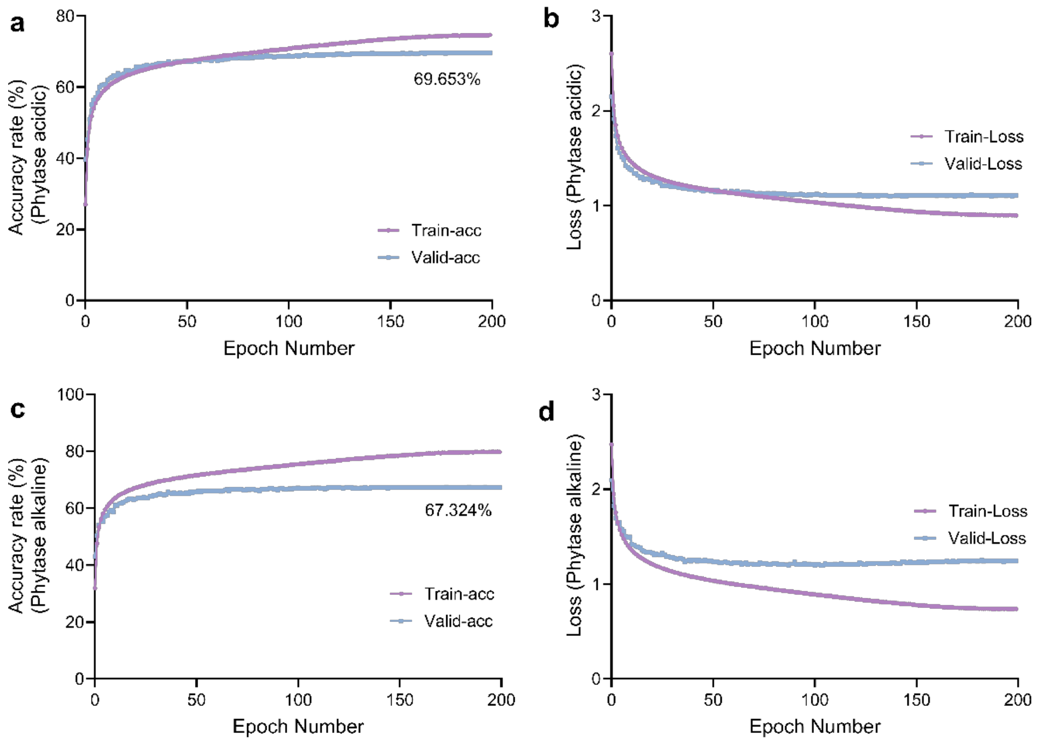

**Fig. S 23 | The verification of CASPE's universality.** (a, b) Results for the verification of CASPE's universality based on phytase protein sequences related to acid stability. (c, d) Results based on phytase protein sequences related to alkaline stability.

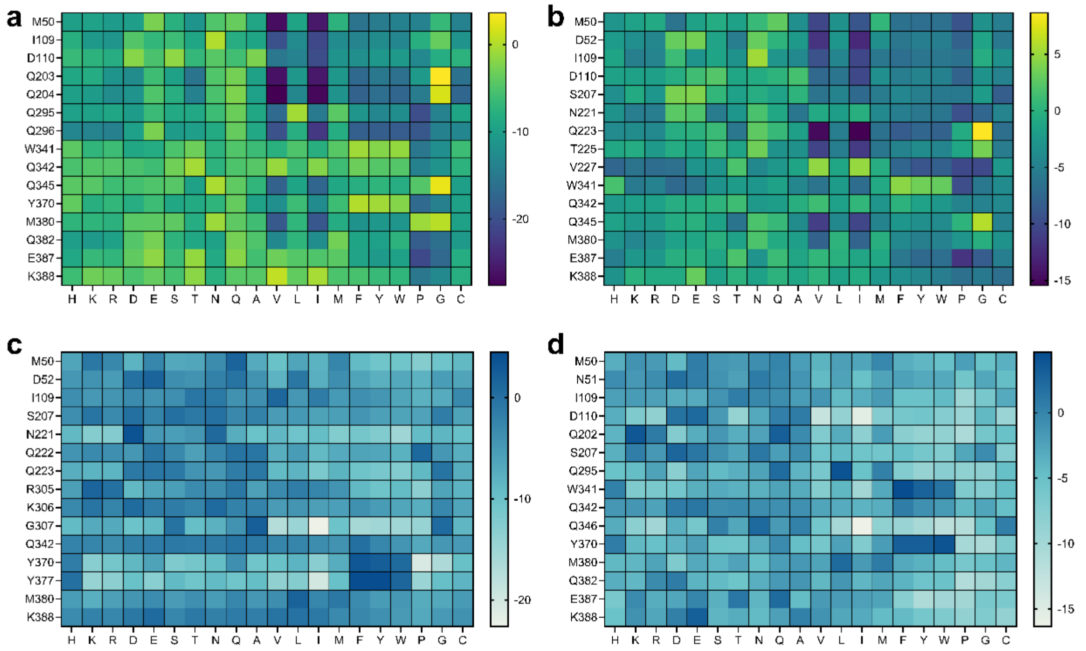

**Fig. S 24 | Heatmaps of single-site mutants selected by CASPEA for phytase.** (a, b) Sites based on phytase related to acid/alkaline stability by CASPEA. (c, d) Sites based on phytase related to acid/alkaline stability by CASPEA-Phytase. (a, c) for acid stability. (b, d) for alkaline stability.

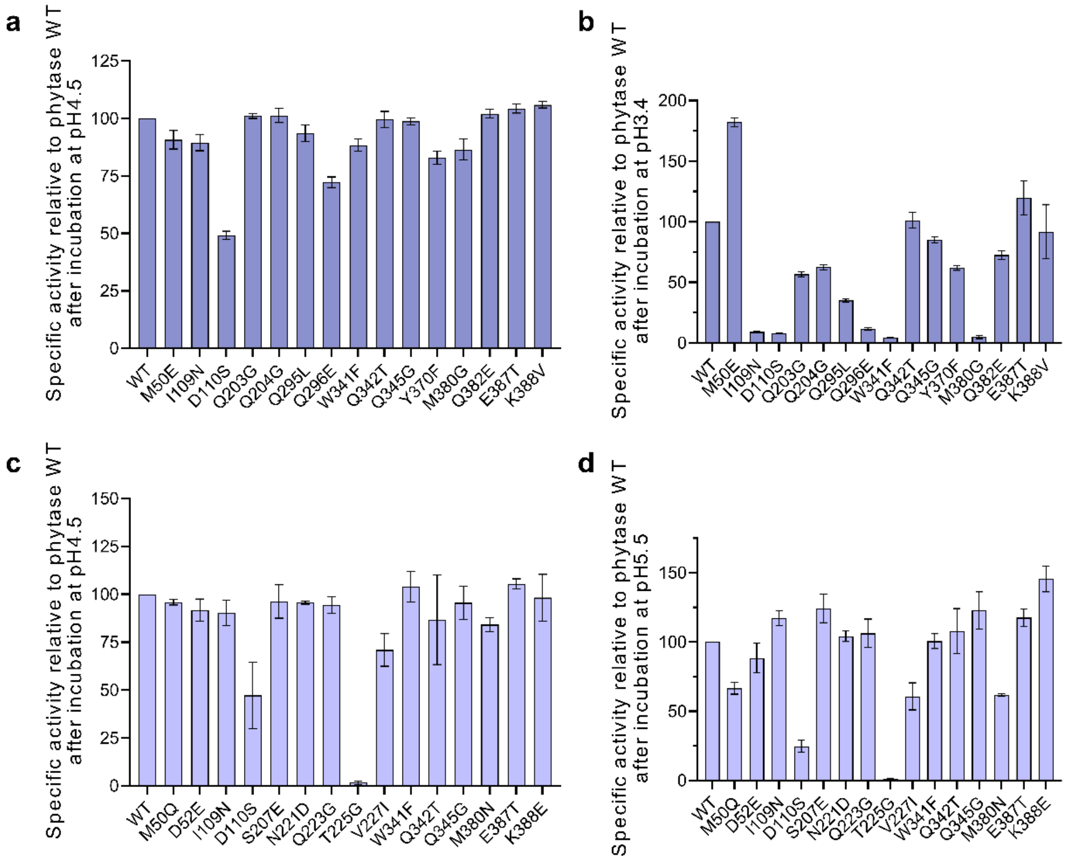

**Fig. S 25 | Specific activity of mutants relative to wild-type, and the mutants were predicted by CASPEA.** (a, b) for acid stability. (c, d) for alkaline stability.

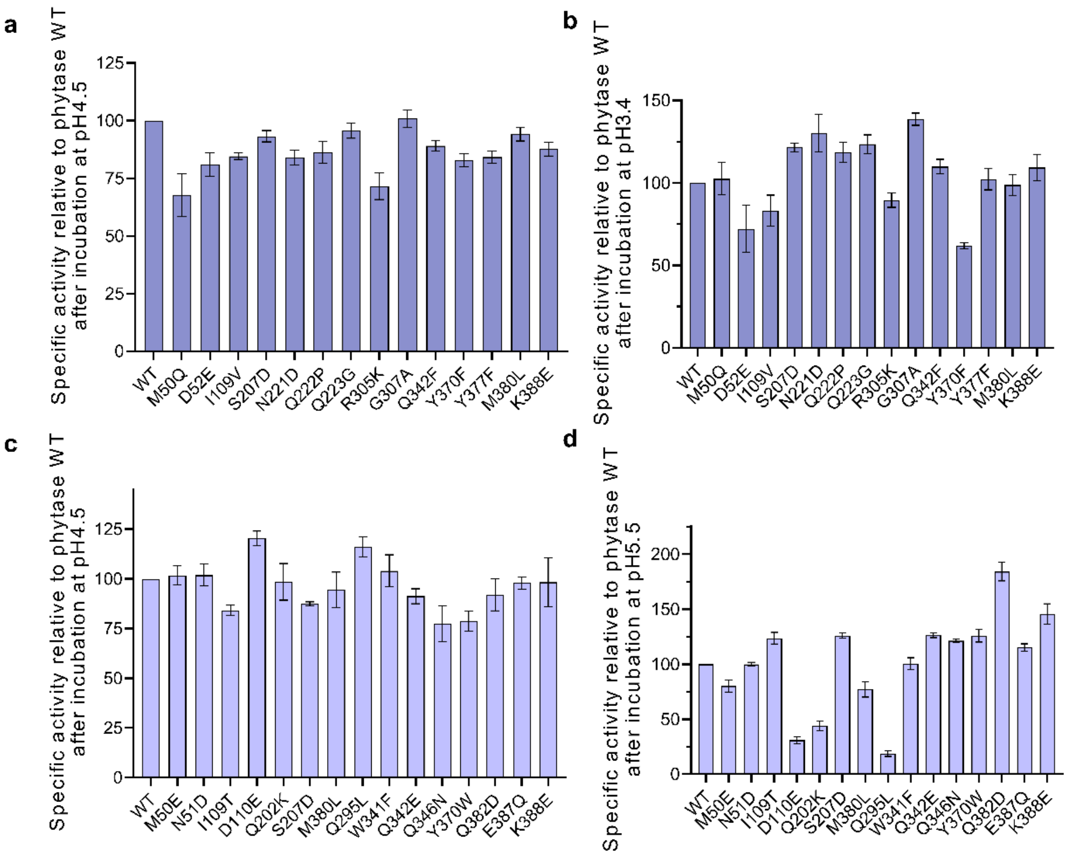

**Fig. S 26 | Specific activity of mutants relative to wild-type, and the mutants were predicted by CASPEA-Phytase.** (a, b) for acid stability. (c, d) for alkaline stability.

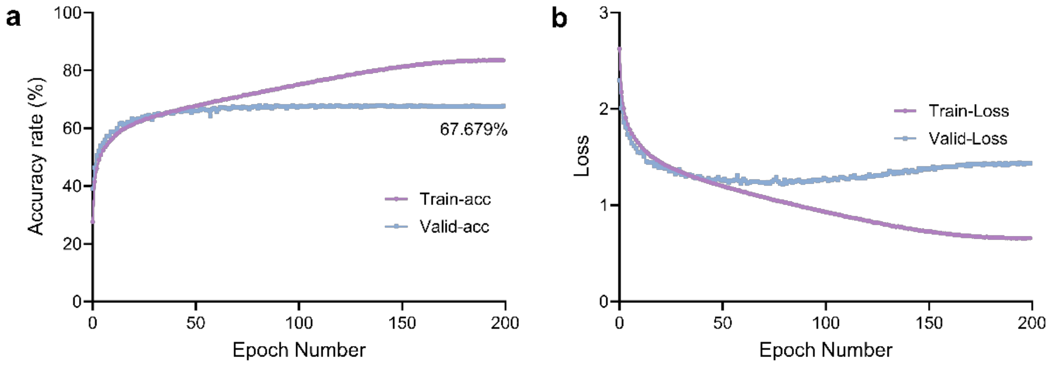

**Fig. S 27 | The verification of CASPE's universality.** (a, b) Results for the verification of CASPE's universality based on phytase protein sequences related to thermostability.

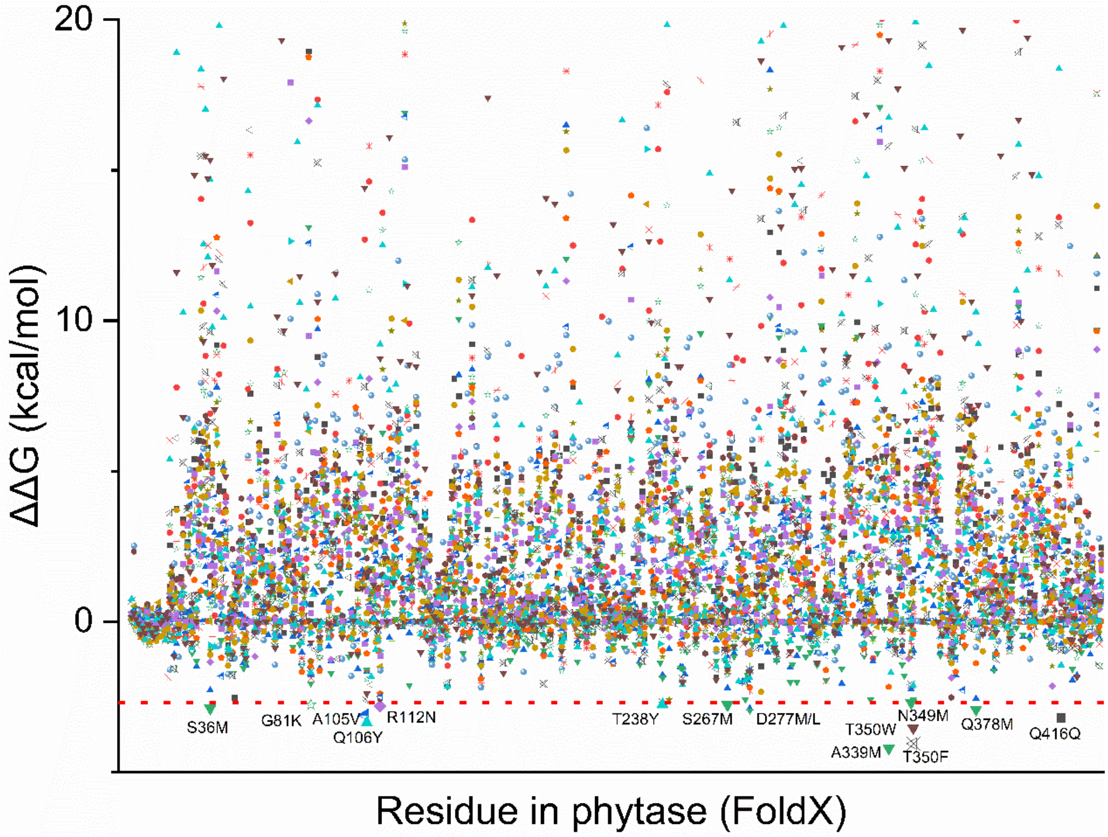

**Fig. S 28 | The mutant sites were screened based on FoldX to enhance the thermostability of phytase, ΔΔG, the smaller the better.**

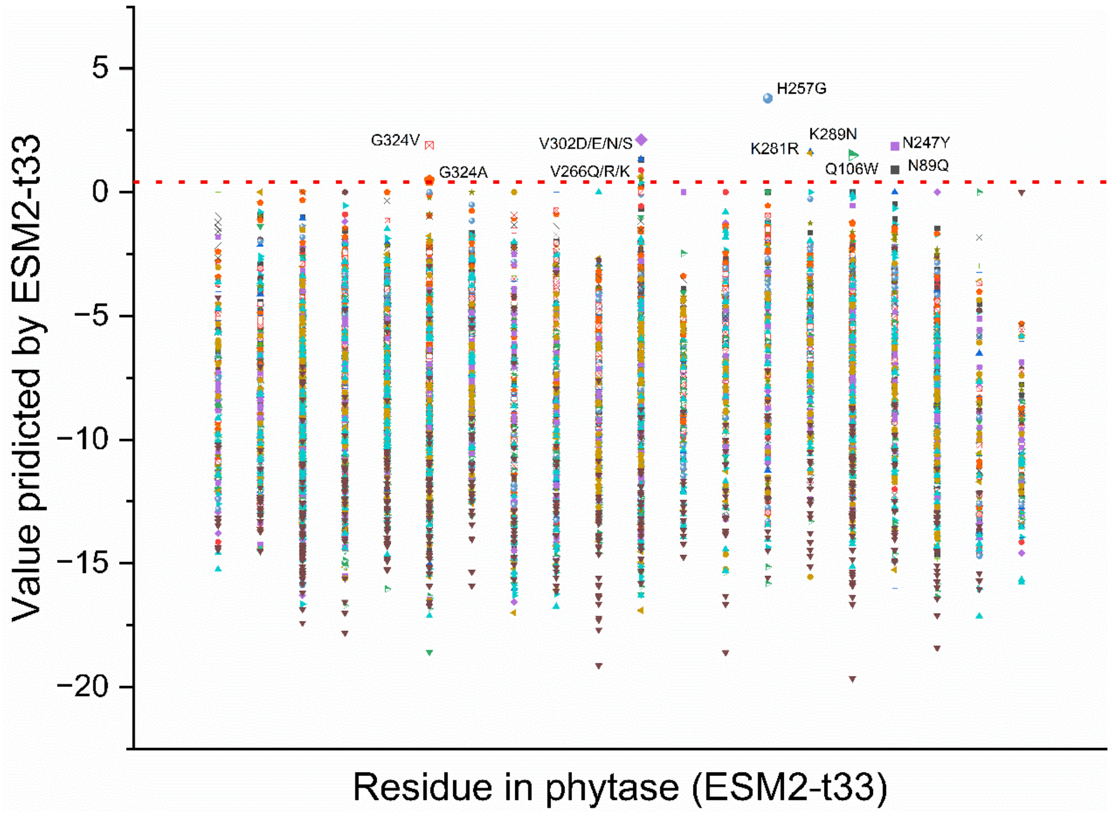

**Fig. S 29 | The mutant sites were screened based on ESM2-t33 to enhance the thermostability of phytase, value, the bigger the better.**

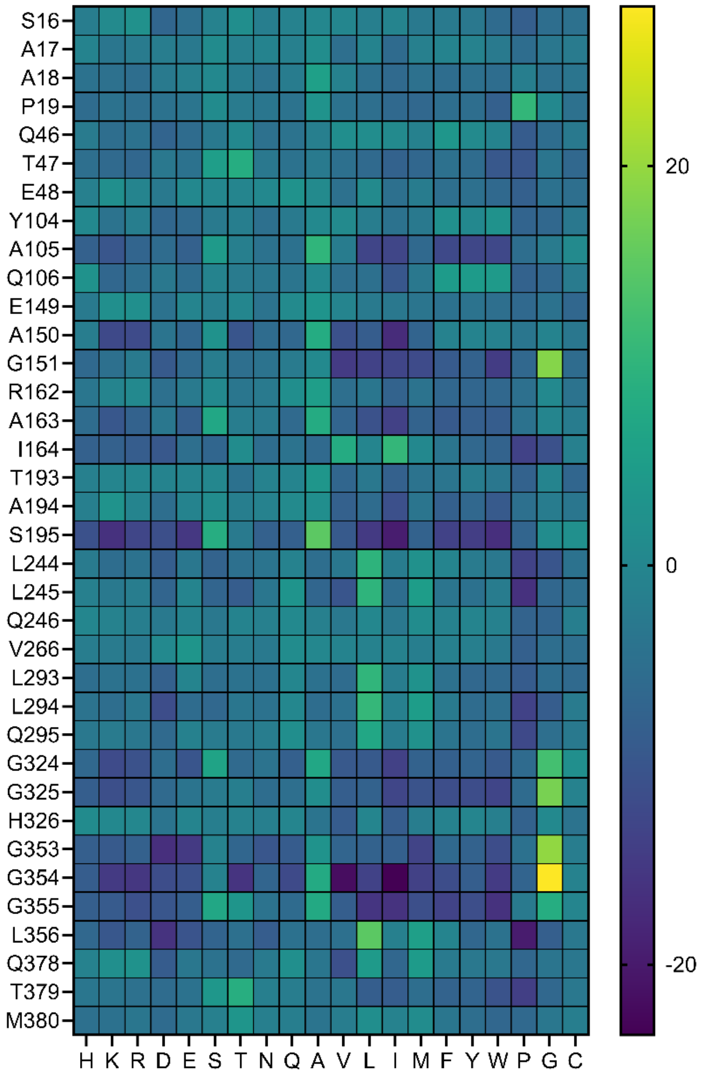

**Fig. S 30 | Heatmaps of single-site mutants selected by CASPET-Phytase for phytase.**

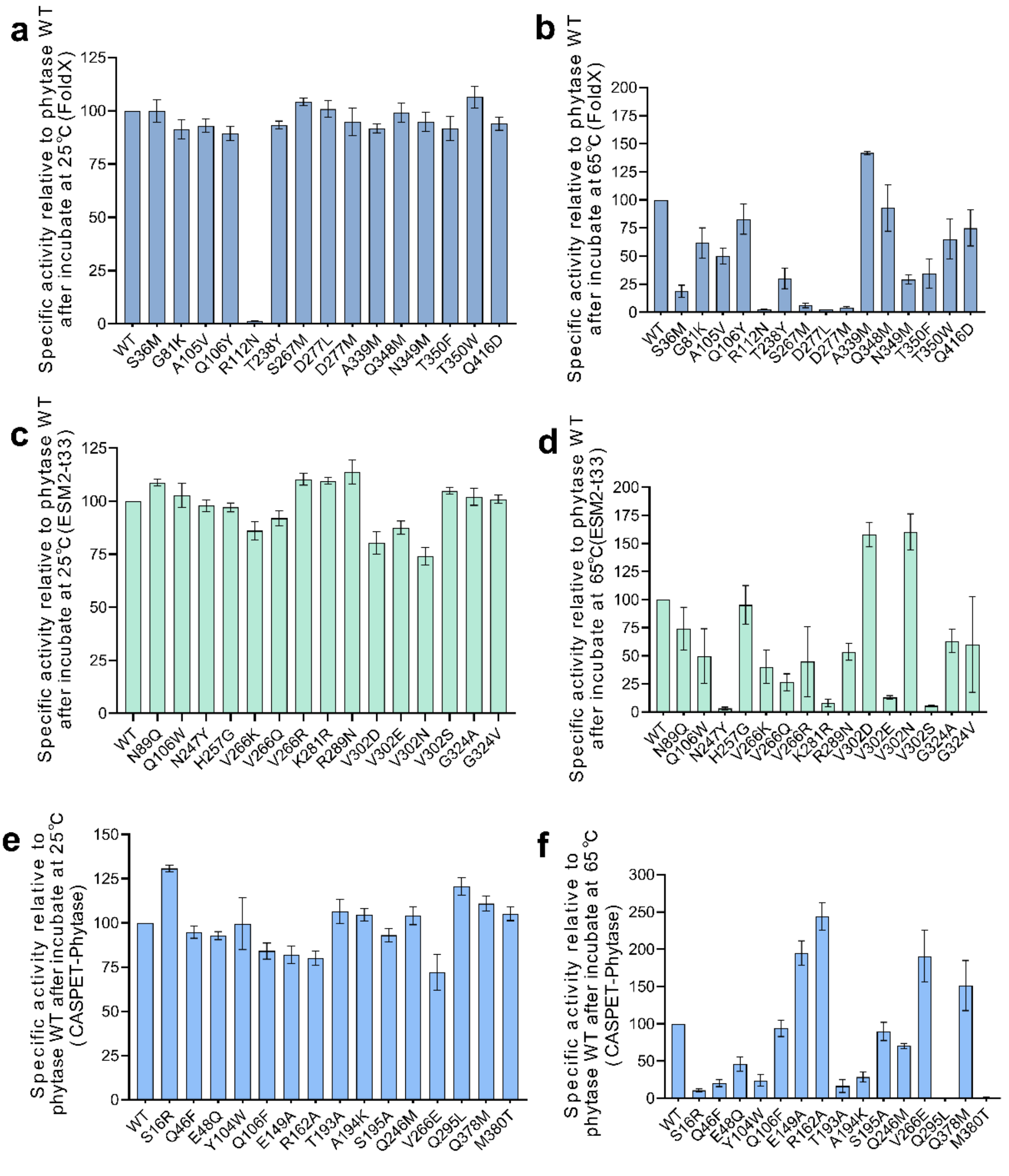

**Fig. S 31 | The relative activities of the mutants obtained by FoldX, ESM2-t33 and CASPET-Phytase at room temperature or high temperature.** (a, c, e) Specific activity relative to wild-type after incubate at 25 °C. (b, d, f) Specific activity relative to wild-type after incubate at 65 °C. (a, b) FoldX. (c, d) ESM2-t33. (e, f) CASPET-Phytase.

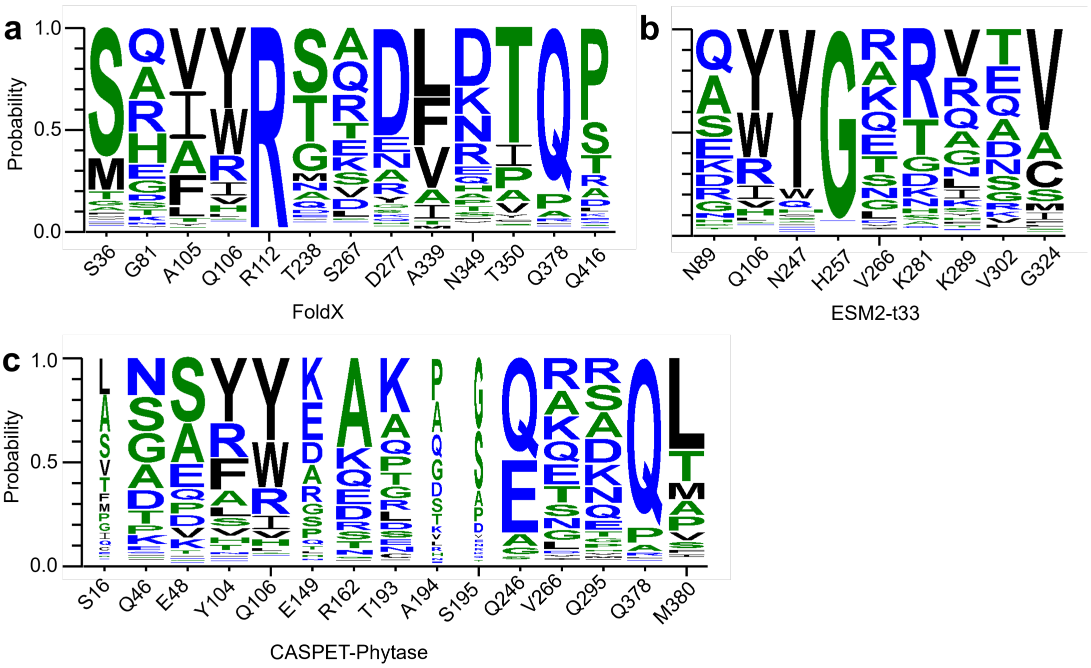

**Fig. S 32 | The conservation analysis of mutation sites selected by FoldX, ESM2-t33 and CASPET-Phytase.** (a) for FoldX, (b) for ESM2-t33 and (c) for CASPET-Phytase.

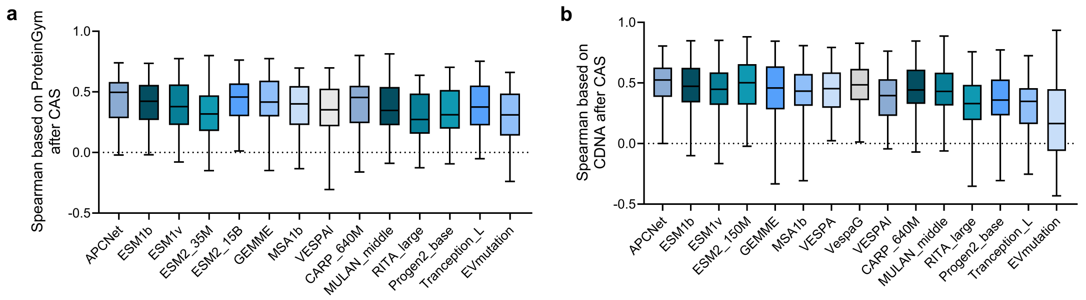

**Fig. S 33 | The results of correlation coefficient analysis based on proteinGym and CDNA**. (a) for proteinGym and (b) for CDNA.

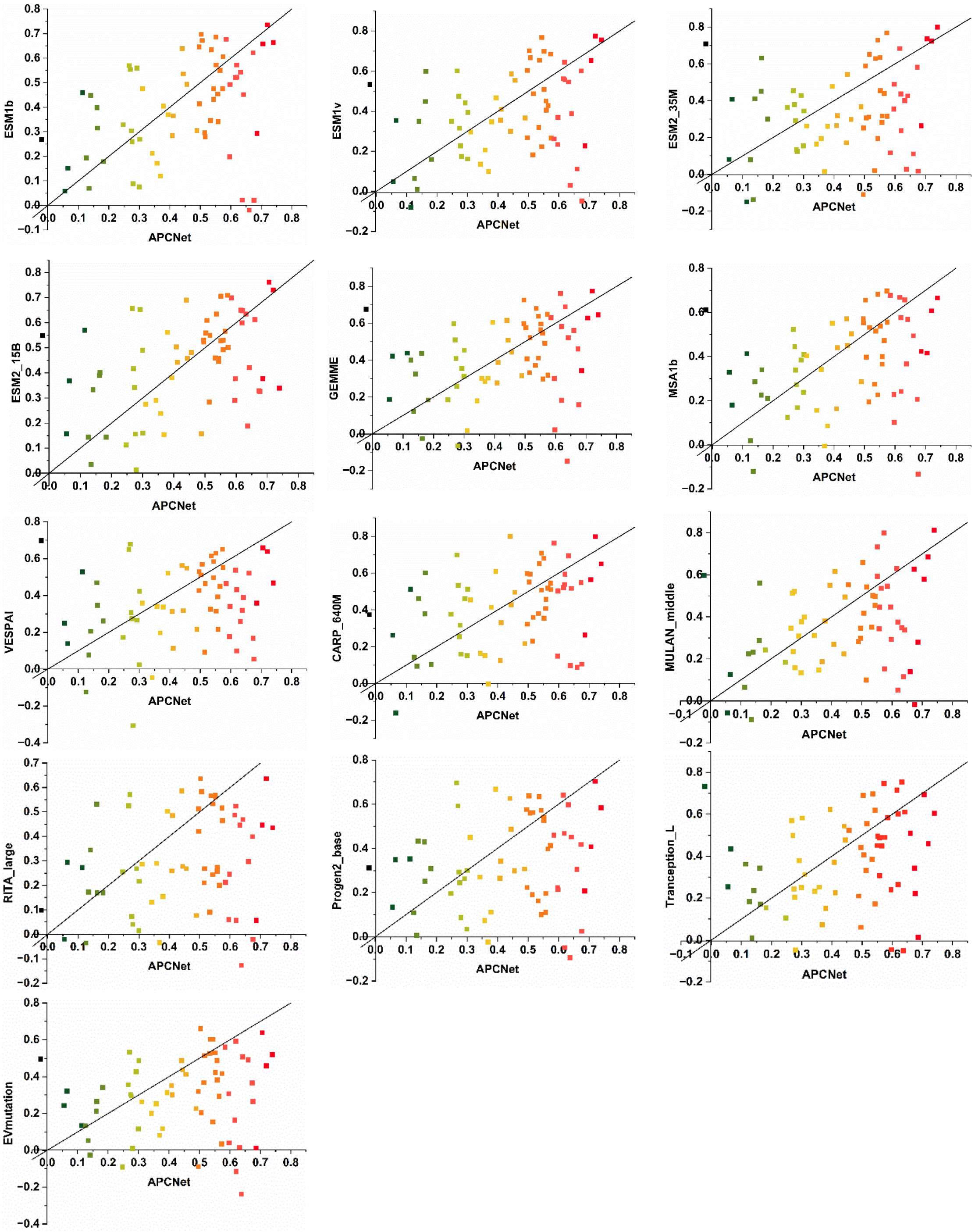

**Fig. S 34 | Details about testing APCNet based on CAS with proteinGym.**

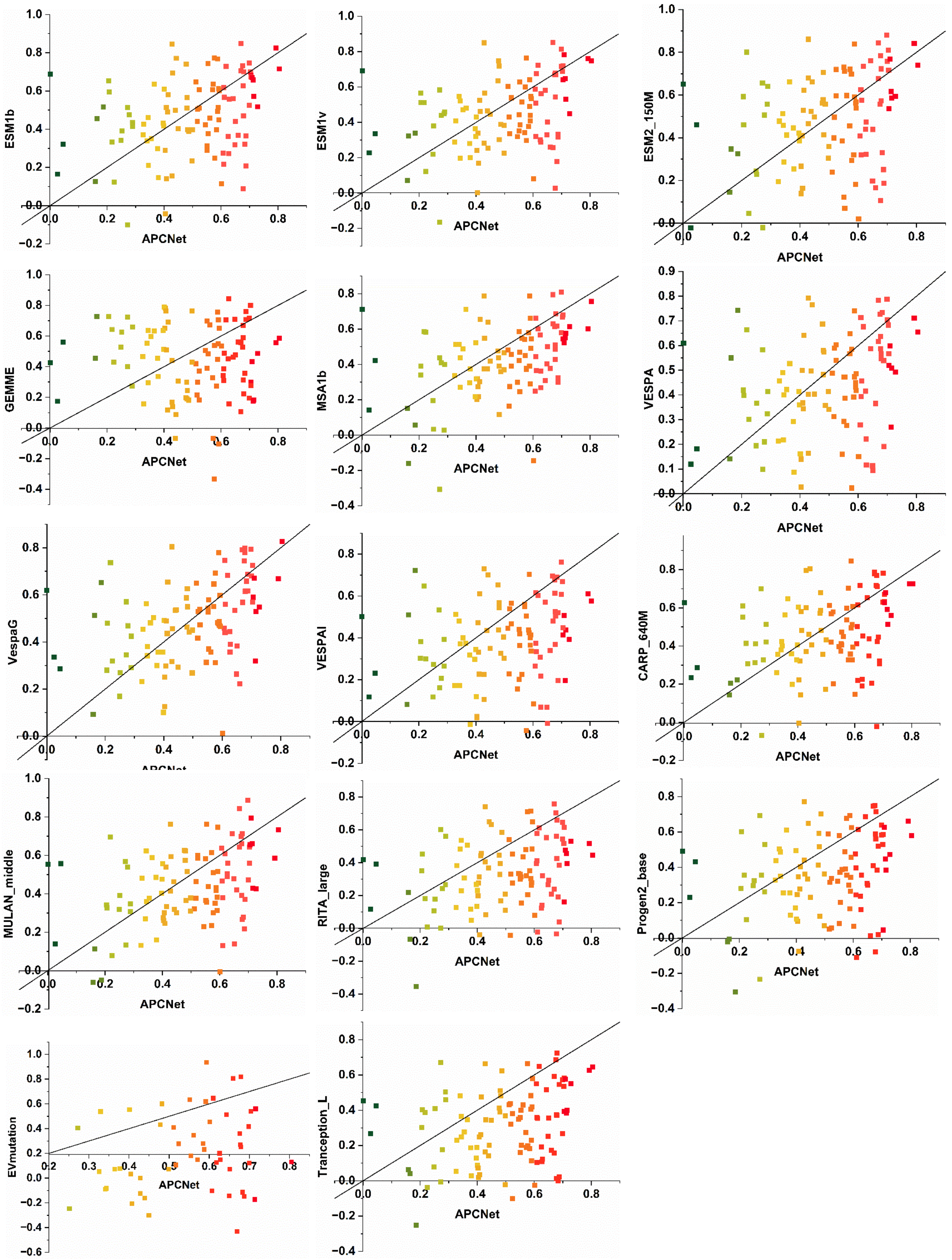

**Fig. S 35 | Details about testing APCNet based on CAS with CDNA.**

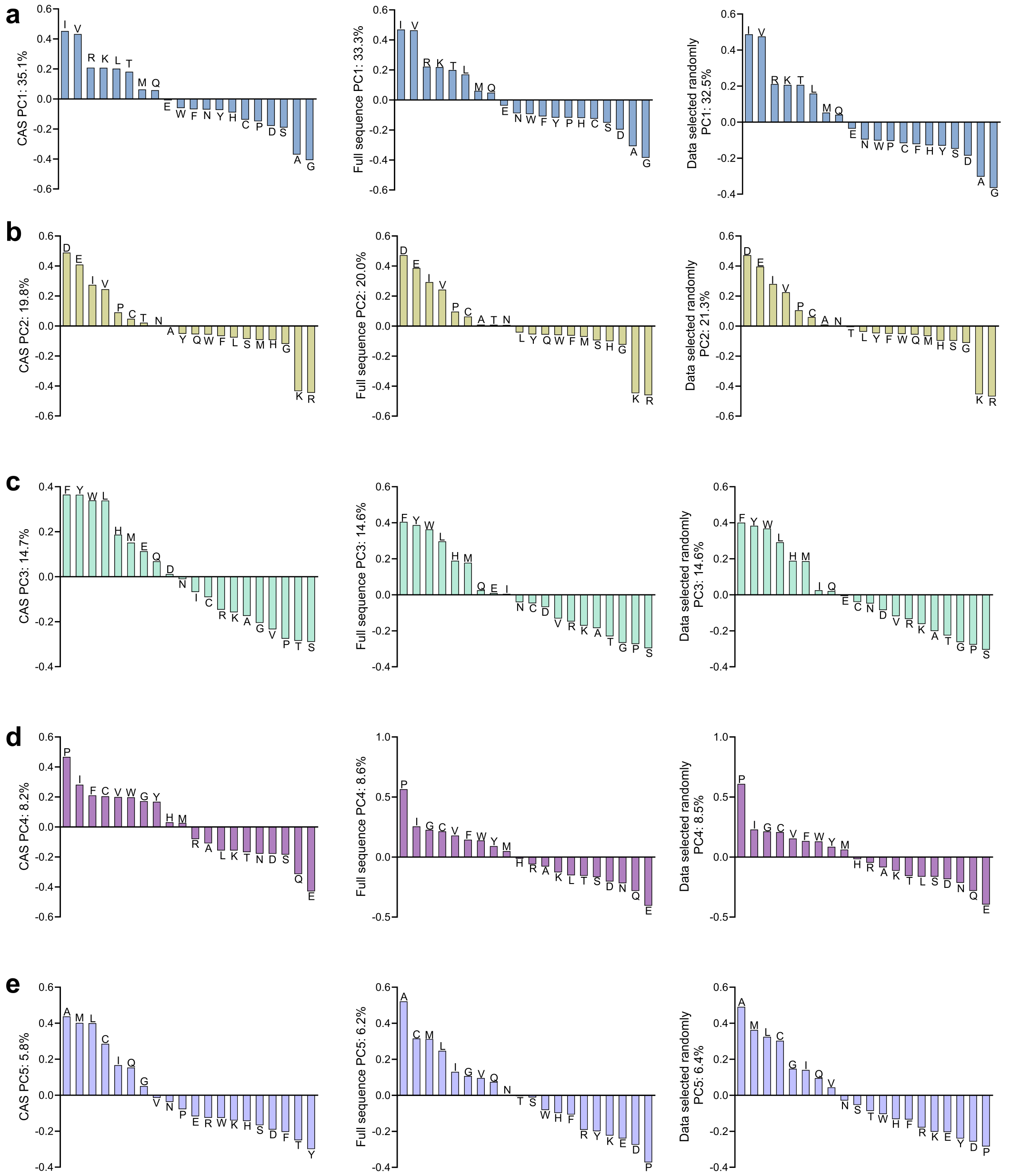

**Fig. S 36 | Results of PCA analysis.** (a-e) PC1-5 of CAS, full sequence and data selected randomly. CAS: 10 sites were selected from each sequence by CAS. Data selected randomly: 10 sites were randomly selected from each sequence.

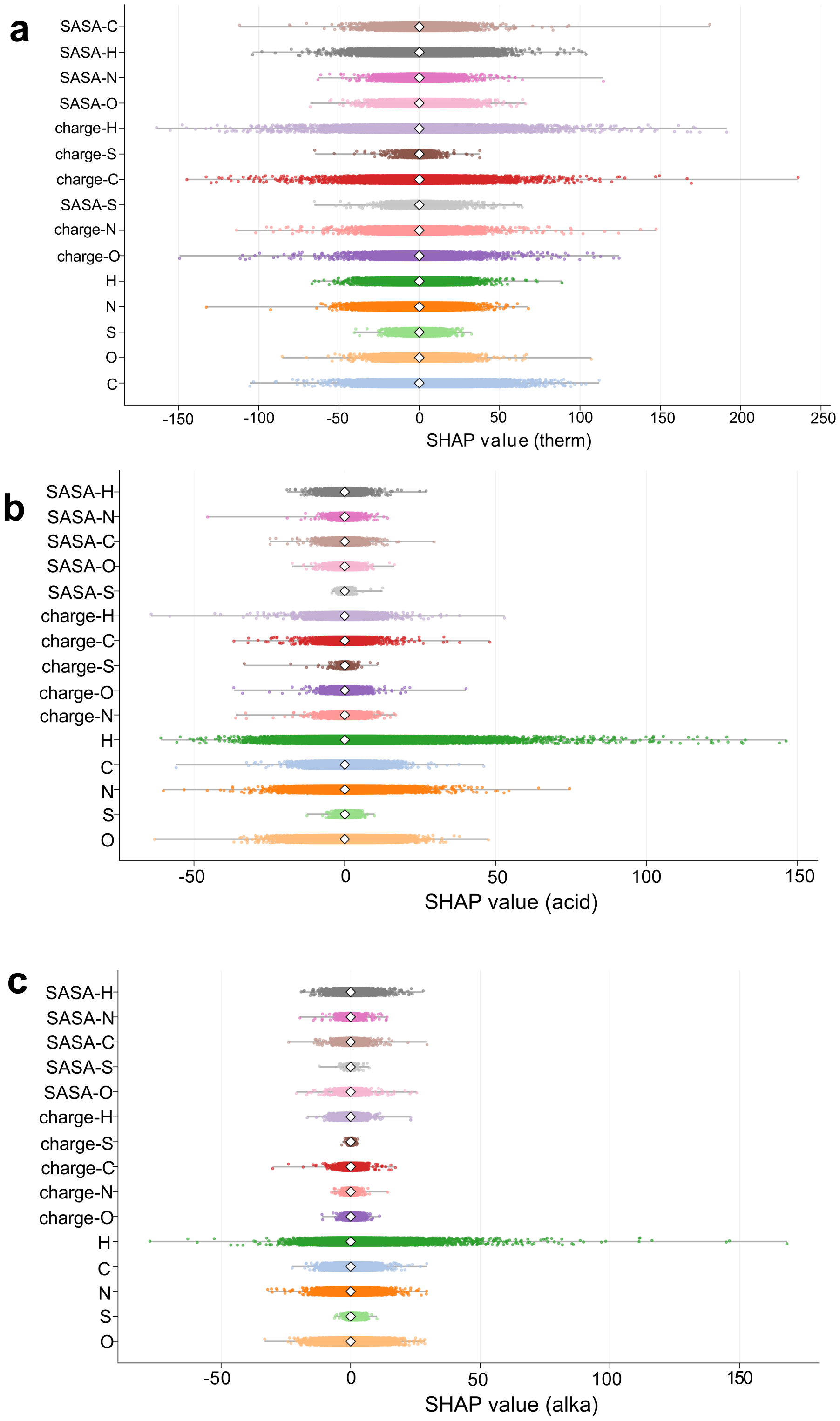

**Fig. S 37 | SHAP distribution of specific values for each feature in all samples of APCNet related to thermostability (a), acid stability (b) and alkaline stability (c).**

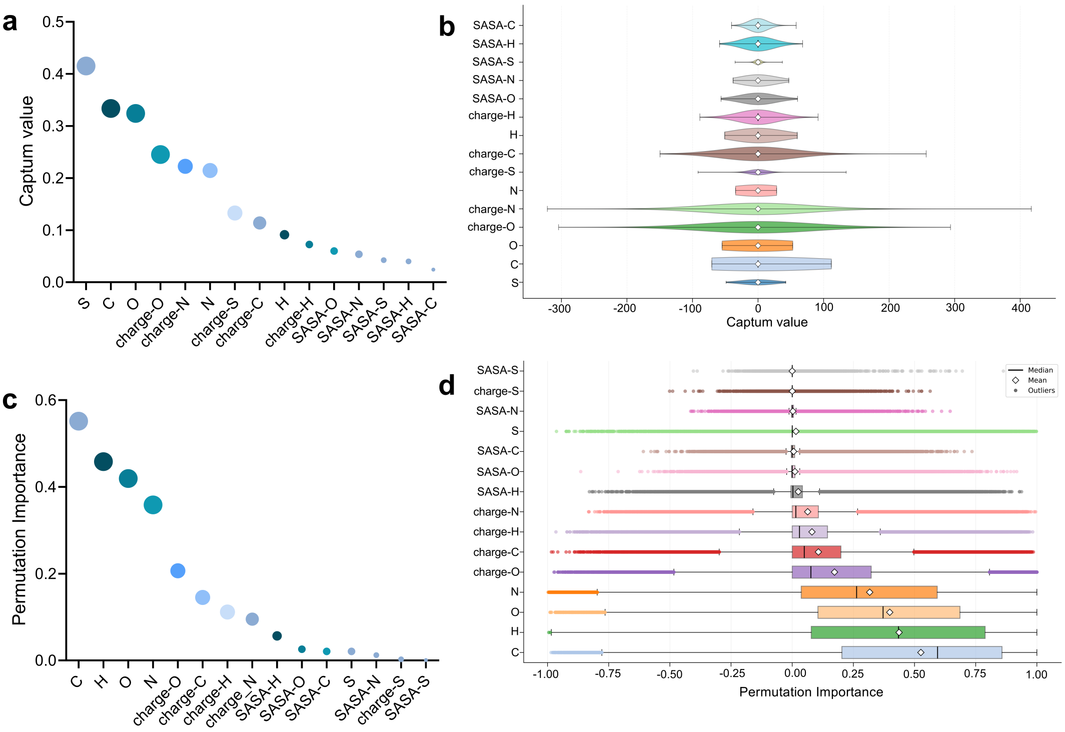

**Fig. S 38 | Interpretability analysis of APCNet related to thermostability.** (a, c) Global analysis of each feature, the influence of each feature on the model is measured by calculating the average absolute value of each feature in all samples. (b, d) The distribution of specific values for each feature in all samples. (a, b) Analysis by Captum. (c, d) Analysis by Permutation Importance.

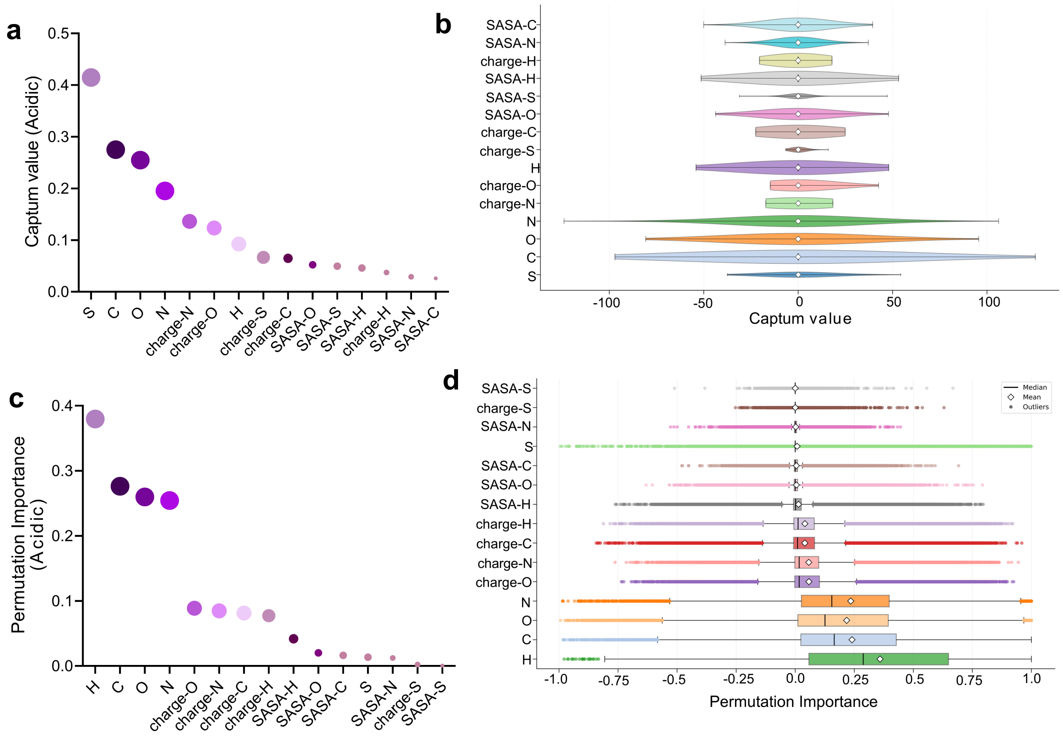

**Fig. S 39 | Interpretability analysis of APCNet related to acid stability.** (a, c) Global analysis of each feature, the influence of each feature on the model is measured by calculating the average absolute value of each feature in all samples. (b, d) The distribution of specific values for each feature in all samples. (a, b) Analysis by Captum. (c, d) Analysis by Permutation Importance.

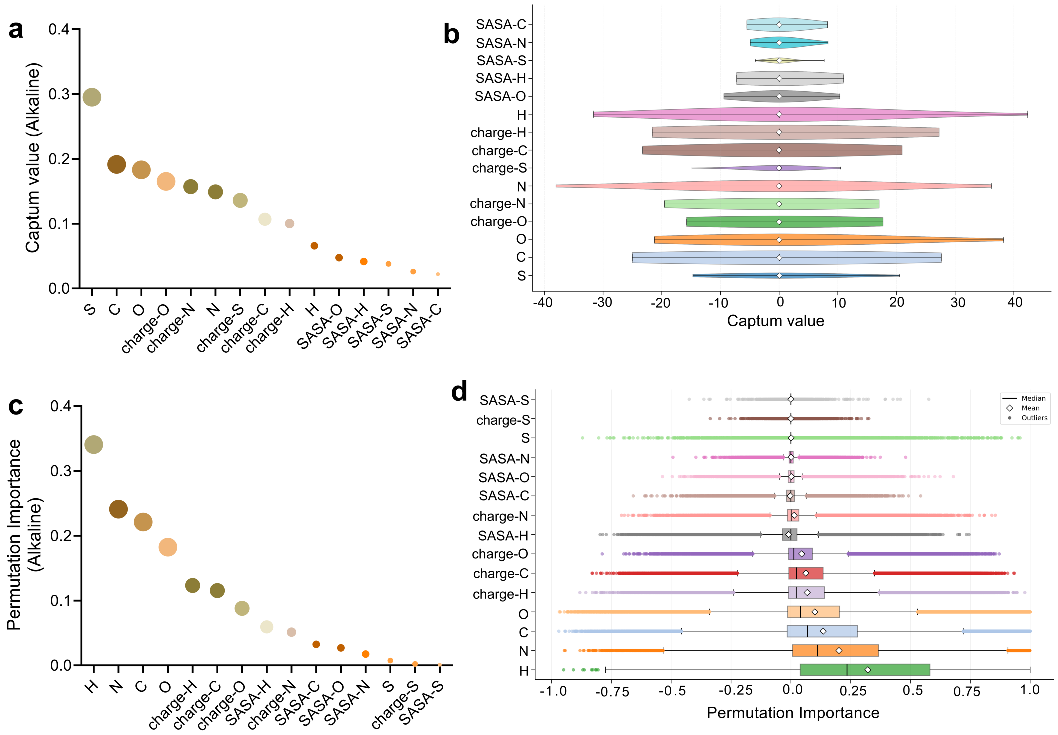

**Fig. S 40 | Interpretability analysis of APCNet related to alkaline stability.** (a, c) Global analysis of each feature, the influence of each feature on the model is measured by calculating the average absolute value of each feature in all samples. (b, d) The distribution of specific values for each feature in all samples. (a, b) Analysis by Captum. (c, d) Analysis by Permutation Importance.

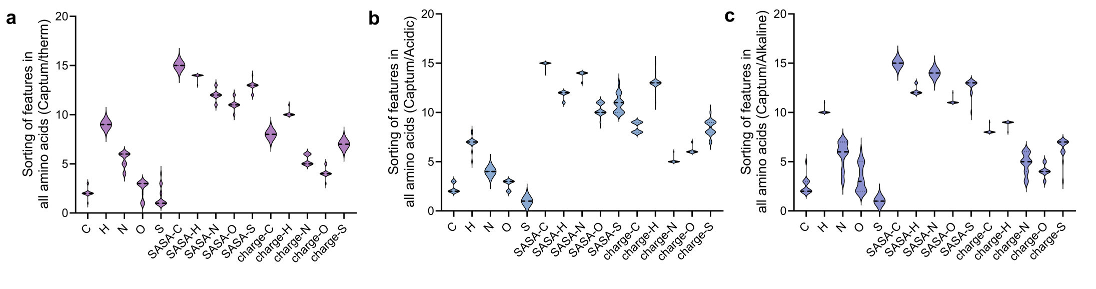

**Fig. S 41 | Count the ranking of each feature among the 20 kinds of amino acids (Captum).** (a) for thermostability, (b) for acid stability and (c) for alkaline stability.

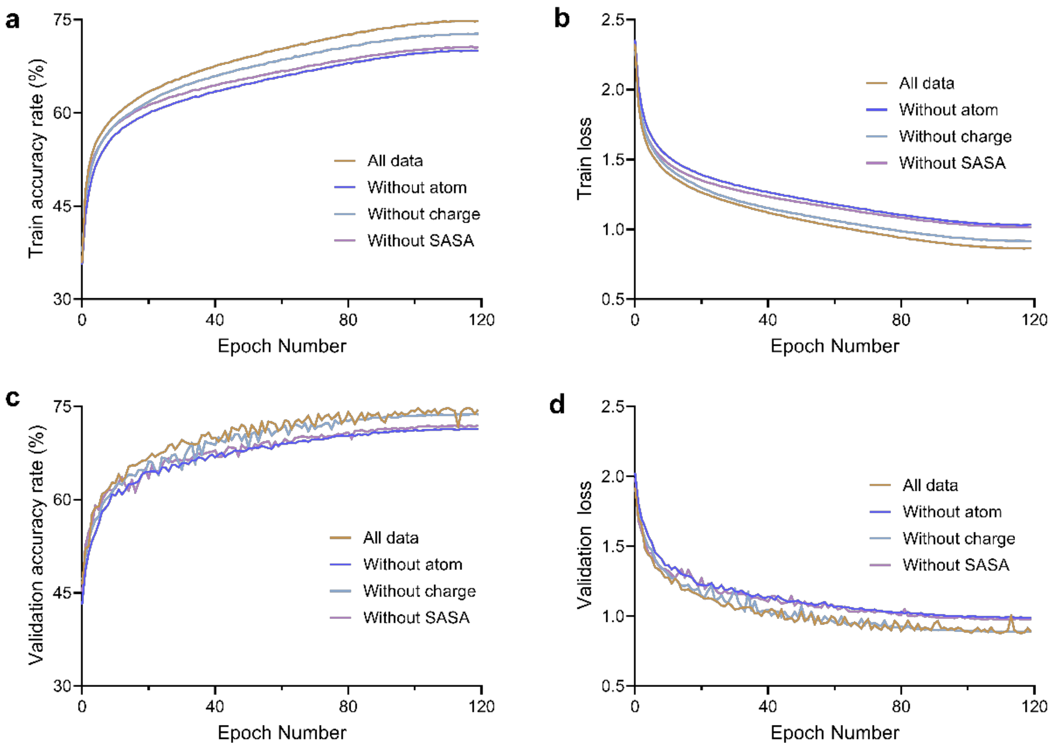

**Fig. S 42 | Ablation experiment based on amino acid microenvironment.** (a, b) Train results of accuracy and loss. (c, d) Valid results of accuracy and loss.

**Table S 1 | Classification model training data gradient setting.**

| | Temperature range (℃) | | | Val-acc |
|---|---|---|---|---|
| **Model 1** | 20-25 | 50-55 | 70-80 | 82.71% |
| **Model 2** | 32-35 | 55-60 | 80-more | 88.50% |
| **Model 3** | Less-20 | 38-45 | 65-70 | 75.66% |
| **Model 4** | 26 | 45-50 | 60-65 | 78.77% |

**Table S 2 | Sequence of CBHI, EG, BG.**

| | Sequence |
|---|---|
| CBHI | QSACTLQSETHPPLTWQKCSSGGTCTQQTGSVVIDANWRWTHAT<br>NSSTNCYDGNTWSSTLCPDNETCAKNCCLDGAAYASTYGVTTSG<br>NSLSIGFVTQSAQKNVGARLYLMASDTTYQEFTLLGNEFSFDVDV<br>SQLPCGLNGALYFVSMDADGGVSKYPTNTAGAKYGTGYCDSQC<br>PRDLKFINGQANVEGWEPSSNNANTGIGGHGSCCSEMDIWEANSI<br>SEALTPHPCTTVGQEICEGDGCGGTYSDNRYGGTCDPDGCDWNP<br>YRLGNTSFYGPGSSFTLDTTKKLTVVTQFETSGAINRYYVQNGVT<br>FQQPNAELGSYSGNELNDDYCTAEEAEFGGSSFSDKGGLTQFKKA<br>TSGGMVLVMSLWDDYYANMLWLDSTYPTNETSSTPGAVRGSCST<br>SSGVPAQVESQSPNAKVTFSNIKFGPIGSTGNPSG |
| EG | ANSKEVKKRASSFEWFGSNESGAEFGSGNIPGVEGTDYTFPNTTAI<br>QILIDAGMNIFRVPFLMERMIPTEMTGSLDTAYFEGYSEVINYITGK<br>GAHAVVDPHNFGRYYGTPISSTSDFQTFWSTLASQFKSNDLVIFDT<br>NNEYHDMDESVVVALNQAAIDGIRDAGATTQYIFVEGNAYSGAW<br>TWTTYNTAMVNLTDPSDLIVYEMHQYLDSDGSGTSDQCVSSTVG<br>QERVVDATTWLQSNGKLGILGEFAGGANSVCEEAVEGMLDYLAE<br>NSDVWLGASWWSAGPWWQDYIYSMEPPNGIAYESYLSILETYF |
| BG | MTDHKALAARFPGDFLFGVATASFQIEGATKVDGRKPSIWDAFCN<br>MPGHVFGRHNGDVACDHYNRWEDDLDLIKEMGVEAYRFSIAWP<br>RIIPDGFGPINEKGLDFYDRLVDGCKARGIKTYATLYHWDLPLTLM<br>GDGGWASRSTAHAFQRYAKTVMARLGDRLDAVATFNEPWCAVW<br>LSHLYGIHAPGERNMEAALAAMHHINLAHGFGVEASRHVAPKVP<br>VGLVLNAHSVIPASDSDADLKAAERAFQFHNGAFFDPVFKGEYPA<br>EMMEALGSRMPVVEAEDLSIISQKLDWWGLNYYTPMRVADDAT<br>EGAEFPATKQAPAVSDVKTDIGWEVYAPALHSLVETLYERYELPDC<br>YITENGACYNMGVENGEVDDQPRLDYYAEHLGIVADLVKDGYPV<br>RGYFAWSLMDNFEWAEGYRMRFGLVHVDYETQVRTLKNSGKW<br>YSALASGFPKGNHGVMKG |

**Table S 3 | $T_m$ of BG and EG.**

| BG (°C) | | EG (°C) | |
|---|---|---|---|
| **WT** | 55.6±0.1 | **WT** | 83.53±0.06 |
| **H229F** | 54.9±0.0 | **T91I** | 83.43±0.06 |
| **D240N** | 58.53±0.06 | **G92S** | 83.3±0.0 |
| **D306A** | 59±0.0 | **S146N** | 83.93±0.06 |
| **Q319A** | 55.37±0.15 | **A234T** | 84.6±0.0 |
